# The RNA-binding ubiquitin ligase MKRN1 functions in ribosome-associated quality control of poly(A) translation

**DOI:** 10.1101/516005

**Authors:** Andrea Hildebrandt, Mirko Brüggemann, Susan Boerner, Cornelia Rücklé, Jan Bernhard Heidelberger, Annabelle Dold, Anke Busch, Heike Hänel, Andrea Voigt, Stefanie Ebersberger, Ingo Ebersberger, Jean-Yves Roignant, Kathi Zarnack, Julian König, Petra Beli

**Affiliations:** Institute of Molecular Biology (IMB), Ackermannweg 4, 55128 Mainz, Germany; Buchmann Institute of Molecular Life Sciences (BMLS), Goethe University, Max-von-Laue-Str. 15, 60438 Frankfurt am Main, Germany; Department for Applied Bioinformatics, Institute of Cell Biology and Neuroscience, Goethe University, Max-von-Laue-Str. 13, 60438 Frankfurt am Main, Germany; Senckenberg Biodiversity and Climate Research Centre (BiK-F), Georg-Voigt-Straße 14-16, 60325 Frankfurt am Main, Germany

**Author notes:** Corresponding authors: KZ, JK and PB.

**Keywords:** MKRN1, ubiquitylation, RNA binding, ribosome-associated quality control, RQC, poly(A), iCLIP, ubiquitin remnant profiling, translation

## Abstract

Cells have evolved quality control mechanisms to ensure protein homeostasis by detecting and degrading aberrant mRNAs and proteins. A common source of aberrant mRNAs is premature polyadenylation, which can result in non-functional protein products. Translating ribosomes that encounter poly(A) sequences are terminally stalled, followed by ribosome recycling and decay of the truncated nascent polypeptide via the ribosome-associated quality control (RQC). Here, we demonstrate that the conserved RNA-binding E3 ubiquitin ligase Makorin Ring Finger Protein 1 (MKRN1) promotes ribosome stalling at poly(A) sequences during RQC. We show that MKRN1 interacts with the cytoplasmic poly(A)-binding protein (PABP) and is positioned upstream of poly(A) tails in mRNAs. Ubiquitin remnant profiling uncovers PABP and ribosomal protein RPS10, as well as additional translational regulators as main ubiquitylation substrates of MKRN1. We propose that MKRN1 serves as a first line of poly(A) recognition at the mRNA level to prevent production of erroneous proteins, thus maintaining proteome integrity.

## Introduction

During gene expression, quality control pathways monitor each step to detect aberrant mRNAs and proteins. These mechanisms ensure protein homeostasis and are essential to prevent neurodegenerative diseases (Chu et al. 2009). A common source of aberrant mRNAs is premature polyadenylation, often in combination with mis-splicing, which results in truncated non-functional protein products (Kaida et al. 2010). Therefore, mechanisms are in place that recognise such homopolymeric adenosine (poly(A)) sequences and abrogate their translation (Bengtson and Joazeiro 2010).

In eukaryotes, ribosomes that terminally stall for diverse reasons during translation are detected by the ribosome-associated quality control (RQC) (reviewed in Brandman and Hegde 2016; Joazeiro 2017). Upon splitting of the 60S and 40S ribosomal subunits, the RQC complex assembles on the stalled 60S subunit to initiate the release and rapid degradation of the truncated tRNA-bound polypeptide. The E3 ubiquitin ligase Listerin (LTN1) modifies the truncated polypeptide with K48-linked ubiquitin chains to target it for degradation in a p97-dependent manner through the proteasome (Bengtson and Joazeiro 2010; Brandman et al. 2012; Verma et al. 2013). Whereas peptide release and ribosome recycling by the RQC complex are relatively well understood, less is known about the mechanisms that promote poly(A) recognition and initial ribosome stalling.

Several recent studies demonstrated a role for the RNA-binding E3 ubiquitin ligase ZNF598 in initiating RQC for prematurely polyadenylated mRNAs (Garzia et al. 2017; Juszkiewicz and Hegde 2017; Sundaramoorthy et al. 2017). It was suggested that ZNF598 senses the translation of poly(A) segments through binding the cognate lysine tRNAs (Garzia et al. 2017). In addition, ZNF598 recognises the collided di-ribosome structure that arises when a trailing ribosome encounters a slower leading ribosome (Juszkiewicz et al. 2018). This is followed by site-specific, regulatory ubiquitylation of the 40S ribosomal proteins RPS10 and RPS20 by ZNF598. In addition to ZNF598, the 40S ribosomal subunit-associated protein RACK1 was shown to regulate ubiquitylation of RPS2 and RPS3 upstream of ribosomal rescue (Sundaramoorthy et al. 2017).

Makorin Ring Finger Protein 1 (MKRN1) belongs to a family of evolutionary conserved RNA-binding E3 ubiquitin ligases. Up to four paralogs exist in vertebrates (MKRN1-4), which combine a RING domain with one or more CCCH zinc finger domains (Gray et al. 2000; Böhne et al. 2010) (**Supplemental Fig. 1A**). MKRN1 has been implicated in the regulation of telomere length, RNA polymerase II transcription and the turnover of tumour suppressor protein p53 and cell cycle regulator p21 (Kim et al. 2005; Omwancha et al. 2006; Lee et al. 2009; Salvatico et al. 2010), but its RNA-related functions remain poorly understood. A study in mouse embryonic stem cells (mESC) reported its interaction with hundreds of mRNAs as well as multiple RNA-binding proteins (RBPs), including the cytoplasmic poly(A)-binding protein (PABP) PABPC1, IGF2BP1 and ELAVL1 (Cassar et al. 2015). The interaction with PABP was further corroborated in human HEK293 cells (Miroci et al. 2012). The same study demonstrated that a shortened isoform of MKRN1 controls local translation via its PABP-interacting motif 2 (PAM2 motif) in rat neurons (Miroci et al. 2012). In line with a role in translation, MKRN1 was found in association with ribosomes, from which it could be released together with PABP and other proteins by RNase digestion (Simsek et al. 2017). Nevertheless, the RNA binding specificity and functional role of MKRN1 in human cells remained largely elusive.

Here, we introduce MKRN1 as a novel factor in RQC. We propose that MKRN1 is recruited to A-rich sequences in mRNAs in a PABP-dependent manner, where it acts as a first line of defence against poly(A) translation. MKRN1 depletion abrogates ribosome stalling in reporter assays, accompanied by reduced ubiquitylation of RQC-related proteins. We therefore hypothesise that MKRN1 allows recognition of poly(A) sequences prior to their translation.

## Results

### MKRN1 interacts with PABPC1 and other RBPs

In order to learn about potential functions, we first characterised the protein interaction profile of MKRN1 in HEK293T cells. To this end, we used affinity purification (AP) coupled to stable isotope labelling with amino acids in cell culture (SILAC)-based quantitative mass spectrometry (MS) using GFP-MKRN1^wt^ or GFP as a bait. We identified 53 proteins that were significantly enriched in GFP-MKRN1^wt^ compared to the control APs (false discovery rate [FDR] < 5%, combined ratios of three independent experiments). In line with previous reports (Miroci et al. 2012; Cassar et al. 2015; Hildebrandt et al. 2017), we found the cytoplasmic poly(A)-binding proteins (PABP) PABPC1 and PABPC4 among the highly enriched MKRN1 interactors (z-score > 4, corrected *P* values = 7.18e-10 and 6.16e-16, respectively) (**Fig. 1A**, **Supplemental Fig. S2A**, and **Supplemental Table S1**). Moreover, we detected 14 ribosomal proteins as well as four proteins that were previously shown to co-purify with ribosomes (Simsek et al. 2017), including IGF2BP1, LARP1, UPF1, and ELAVL1 (**Fig. 1A**). Consistently, “translation” was among the significantly enriched Gene Ontology (GO) terms for the MKRN1 interaction partners (Biological Process [BP], **Supplemental Fig. S2B**). Almost all interactors were previously found in association with polyadenylated transcripts (50 out of 53 proteins have been annotated with the GO term “poly(A) RNA binding”, Molecular Function [MF], **Supplemental Fig. S2B**). We confirmed the MS results in reciprocal AP experiments with GFP-tagged PABPC1, ELAVL1, and IGF2BP1 as baits followed by Western blot for endogenous MKRN1 (**Supplemental Fig. S2C**). All detected interactions persisted in the presence of RNases (RNase A and T1), demonstrating that MKRN1 interacts with these proteins in an RNA-independent manner (**Supplemental Fig. S2C**). Together, these observations suggest that MKRN1 is part of a larger mRNA ribonucleoprotein particle (mRNP) together with PABP and other RBPs. This is further supported by a parallel study on the Mkrn1 ortholog in *Drosophila melanogaster*, which consistently identified pAbp, Larp, Upf1 and Imp (IGF2BP in mammals) as interaction partners (Dold et al, parallel submission; preprint available at bioRxiv, doi: 10.1101/501643).

**Figure 1.**
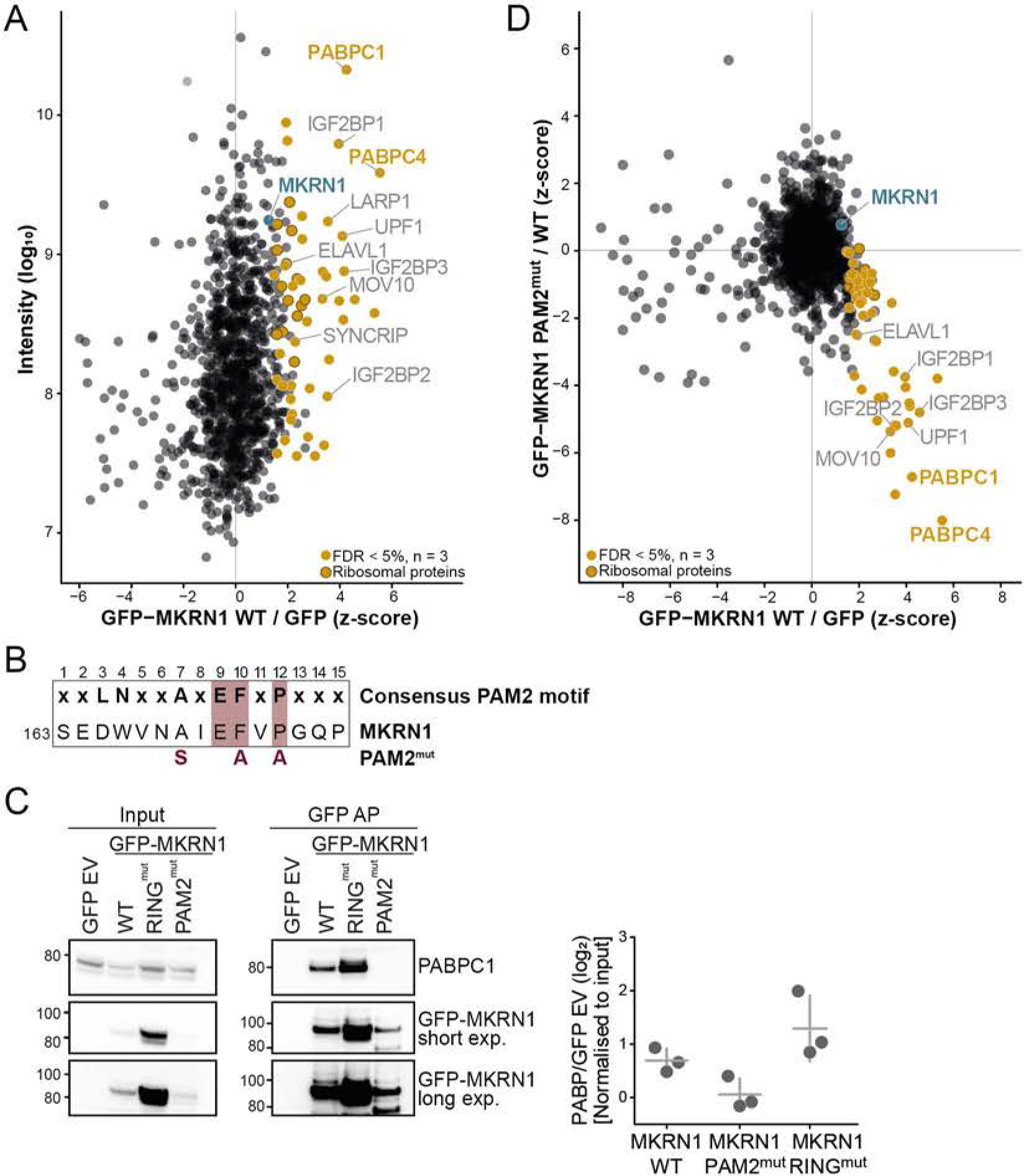
MKRN1 interacts with PABP and other regulators of translation and RNA stability. (*A*) Protein interactome of GFP-MKRN1^wt^ in HEK293T cells analysed by quantitative MS-based proteomics. Combined SILAC ratios (n = 3 replicates) after z-score normalisation are plotted against log_10_-transformed intensities. 1,100 protein groups were quantified in at least two out of three replicate experiments. MKRN1 and significant interactors are highlighted (FDR < 5%). (*B*) A PAM2 motif similar to the previously reported consensus (shown on top; **Supplemental Fig. S1B**) (Albrecht and Lengauer 2004) is present in MKRN1 (first amino acid position indicated on the left). Introduced mutations in MKRN1^PAM2mut^ are indicated in red below. Relevant positions are highlighted (**Supplemental Fig. S1B**). (*C*) Endogenous PABP interacts with MKRN1^wt^ and MKRN1^RINGmut^, but not with MKRN1^PAM2mut^. Western blots for endogenous PABPC1 and GFP (two exposure times, exp.) after AP of GFP-MKRN1 (wt and mutants). Ratios of PABP signal (normalised to input) in GFP-MKRN APs over control (GFP empty vector, EV) are shown on the right. Replicates 2, 3, and uncropped gel images are shown in **Supplemental Fig. S9A-C**. (*D*) Quantitative comparison of the interactomes of GFP-MKRN1^wt^ and GFP-MKRN1^PAM2mut^ shows that PABP and several other interactors are lost upon PAM2 mutation. Combined ratios of three replicates are shown in a scatter plot. Only proteins detected in at least two out of three replicates are shown. MKRN1^wt^ significant interactors (from A) are highlighted as in (*A*) (FDR < 5% in MKRN1^wt^).

Many proteins interact with PABP via a PABP-interacting motif (PAM2) motif, which specifically binds to the MLLE domain present almost exclusively in PABP (Deo et al. 2001; Kozlov et al. 2010). Accordingly, a previous study demonstrated that MKRN1 associates with PABP via a PAM2 motif at amino acid positions 161-193 (Miroci et al. 2012). In support of a putative functional relevance, a phylogenetic analysis illustrated that the presence and positioning of the PAM2 motif are preserved in MKRN1 orthologs across metazoans (**Supplemental Fig. S1A,B**). AP of a MKRN1 variant with point mutations in the PAM2 motif (GFP-MKRN1^PAM2mut^) (Pohlmann et al. 2015) no longer recovered PABPC1 and PABPC4 (**Fig. 1B-D** and **Supplemental Table S1**). For comparison, we also tested a previously described point mutation in the RING domain that abolishes the E3 ubiquitin ligase function (ligase-dead, GFP-MKRN1^RINGmut^) (Kim et al. 2005). This mutation did not impair the interaction of MKRN1 with PABPC1, but led to a slight increase, possibly due to stabilisation of MKRN1 and/or PABPC1 (**Fig. 1C, Supplemental Fig. S7,** and **Supplemental Table S1**). Surprisingly, MKRN1^PAM2mut^ lost interaction not only with PABPC1 and PABPC4, but also with several other identified proteins (**Fig. 1D**), suggesting that MKRN1^PAM2mut^ no longer resided within the mRNPs. These results confirmed that MKRN1 interacts with PABP proteins, and suggested that this association is required for mRNP formation.

### MKRN1 binds to poly(A) tails and at internal A-stretches

In order to characterise the RNA-binding behaviour of human MKRN1 *in vivo*, we performed individual-nucleotide resolution UV crosslinking and immunoprecipitation (iCLIP) (König et al. 2010) in combination with 4-thiouridine (4SU) labelling to enhance UV crosslinking (Hafner et al. 2010). In three replicate experiments with GFP-tagged MKRN1 (GFP-MKRN1^wt^) expressed in HEK293T cells, we identified more than 4,6 million unique crosslink events, cumulating into 7,331 MKRN1 binding sites (see Materials and methods; **Supplemental Table S2**). These were further ranked according to the strength of MKRN1 binding, which was estimated from the enrichment of crosslink events within a binding site relative to its local surrounding, which served as a proxy for transcript abundance (“signal-over-background”, SOB; see Materials and Methods) (Sutandy et al. 2018). SOB values were highly reproducible between replicates (Pearson correlation coefficients *r* > 0.72, **Supplemental Fig. S3**).

Across the transcriptome, MKRN1 almost exclusively bound to protein-coding mRNAs with a strong tendency to locate in 3’ UTRs (**Fig. 2A,E**). Binding sites generally harboured uridine-rich tetramers (**Supplemental Fig. S4A**), likely reflecting 4SU-based UV crosslinking (Hafner et al. 2010). Strikingly, the top 20% MKRN1 binding sites were massively enriched in AAAA tetramers (A, adenosine) within 5-50 nucleotides (nt) downstream of the binding sites (**Fig. 2B** and **Supplemental Fig. S4A**). These were situated within A-rich stretches (A-stretches), which ranged from 8-30 nt in length (**Supplemental Fig. S4B**; see Materials and methods). Within 3’ UTRs, 30% (1,848 out of 6,165) of MKRN1 binding sites resided immediately upstream of an A-stretch (**Fig. 2C,E**) and longer A-stretches associated with stronger MKRN1 binding (**Supplemental Fig. S4C,D**).Intriguingly, we detected a requirement for a continuous run of at least 8 A’s to confer strong MKRN1 binding (**Fig. 2D**), which precisely matched the RNA footprint of PABP (Webster et al. 2018). Since PABP was previously reported to also bind within 3’ UTRs (Bag 2001; Lyabin et al. 2011; Kini et al. 2016), these observations indicated that MKRN1 binds together with PABP to mRNAs.

**Figure 2.**
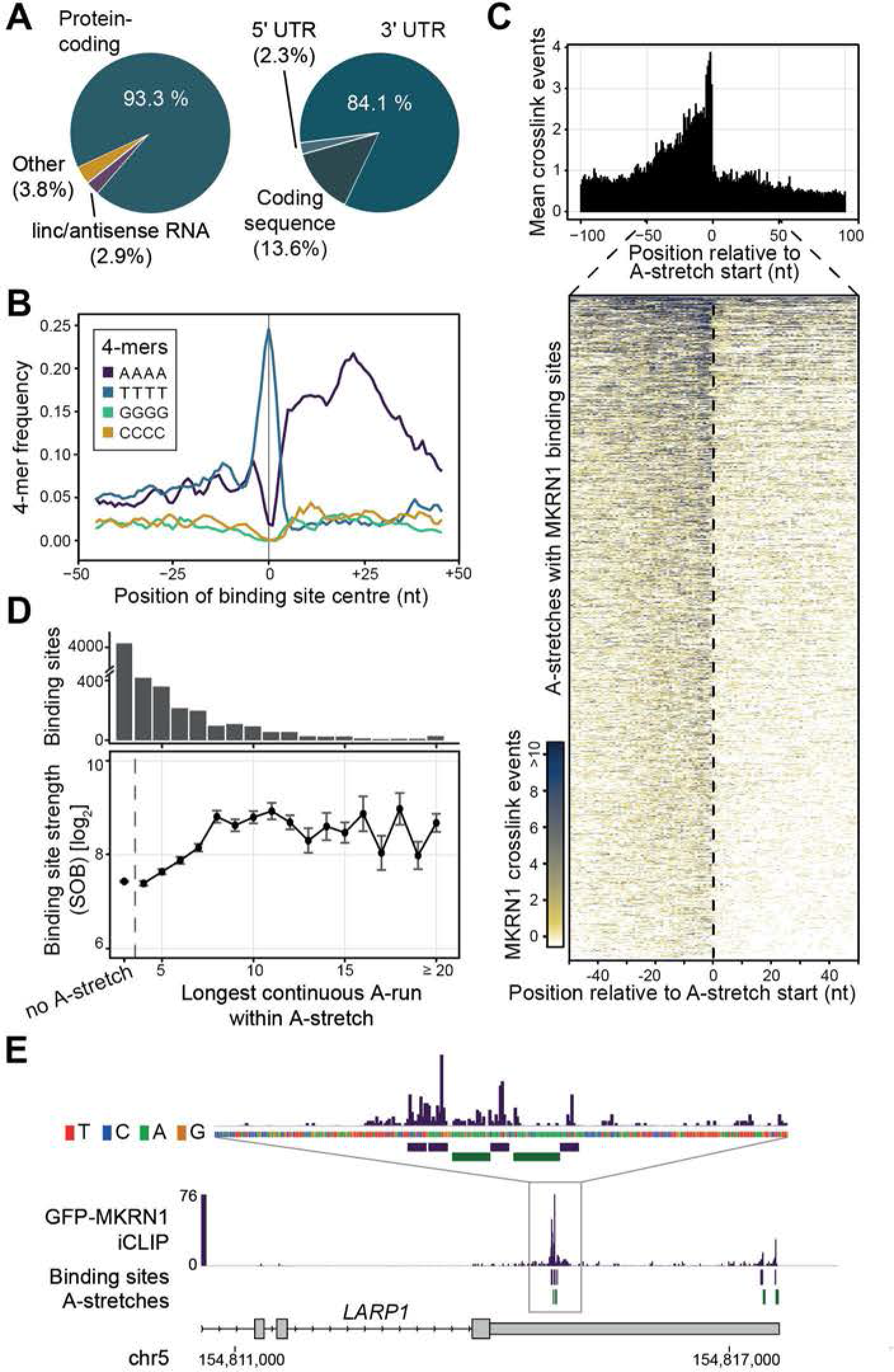
MKRN1 binds upstream of A-stretches in 3’ UTRs. (*A*) MKRN1 predominantly binds in the 3’ UTR of protein-coding genes. Piecharts summarising the distribution of MKRN1 binding sites to different RNA biotypes (7,331 binding sites, left) and different regions within protein-coding transcripts (6,913 binding sites, right). (*B*) MKRN1 binding sites display a downstream enrichment of AAAA homopolymers. Frequency per nucleotide (nt) for four homopolymeric 4-mers in a 101-nt window around the midpoints of the top 20% MKRN1 binding sites (according to signal-over-background; see Material and methods). (*C*) MKRN1 crosslink events accumulate upstream of A-stretches. Metaprofile (top) shows the mean crosslink events per nt in a 201-nt window around the start position of 1,412 MKRN1-associated A-stretches in 3’ UTRs. Heatmap visualisation (bottom) displays crosslink events per nt (see colour scale) in a 101-nt window around the MKRN1-associated A-stretches. (*D*) MKRN1 binding site strength (signal-over-background, SOB) increases with length of longest continuous run of A’s (LCA) within the A-stretch. Mean and standard deviation of MKRN1 binding sites associated with A-stretches harbouring LCAs of increasing length (x-axis). MKRN1 binding sites without associated A-stretches are shown for comparison on the left. Number of binding sites in each category indicated as barchart above. (*E*) MKRN1 binds upstream of A-stretches in the 3’ UTR of the *LARP1* gene. Genome browser view of GFP-MKRN1 iCLIP data showing crosslink events per nt (merged replicates, turquois) together with binding sites (lilac) and associated A-stretches (dark green).

Prompted by this notion, we analysed the unusually high fraction of unmapped iCLIP reads in the MKRN1 dataset (**Supplemental Table S2**). In accordance with binding of MKRN1 immediately upstream of poly(A) tails, more than 13% of the unmapped reads displayed an increased A-content (**Fig. 3B**), compared to only 2% for an unrelated control RBP (Braun et al. 2018). In addition, the mapped GFP-MKRN1^wt^ crosslink events were enriched upstream of annotated polyadenylation sites, as exemplified in the *SRSF4* gene (**Fig. 2E** and **Fig. 3A,C**). Together, these results support the notion that MKRN1 binds upstream of poly(A) tails, possibly in conjunction with PABP. In order to test whether PABP is required for MKRN1 binding, we performed UV crosslinking experiments with GFP-MKRN1^PAM2mut^, which no longer interacts with PABP (**Fig. 1C,D**). Strikingly, RNA binding of this mutant was globally reduced compared to GFP-MKRN1^wt^ (**Fig. 3D** and **Supplemental Fig. S5**), indicating that PABP might recruit MKRN1 to RNA. In summary, these results strongly imply that MKRN1 binds upstream of poly(A) tails, which could be implemented via its interaction with PABP. In a concordant scenario, it was found that *Drosophila* Mkrn1 bound before an extended A-stretch in the 3’ UTR of *oskar* mRNA, and that this binding was significantly reduced upon depletion of pAbp (Dold et al., parallel submission; preprint available at bioRxiv, doi: 10.1101/501643).

**Figure 3.**
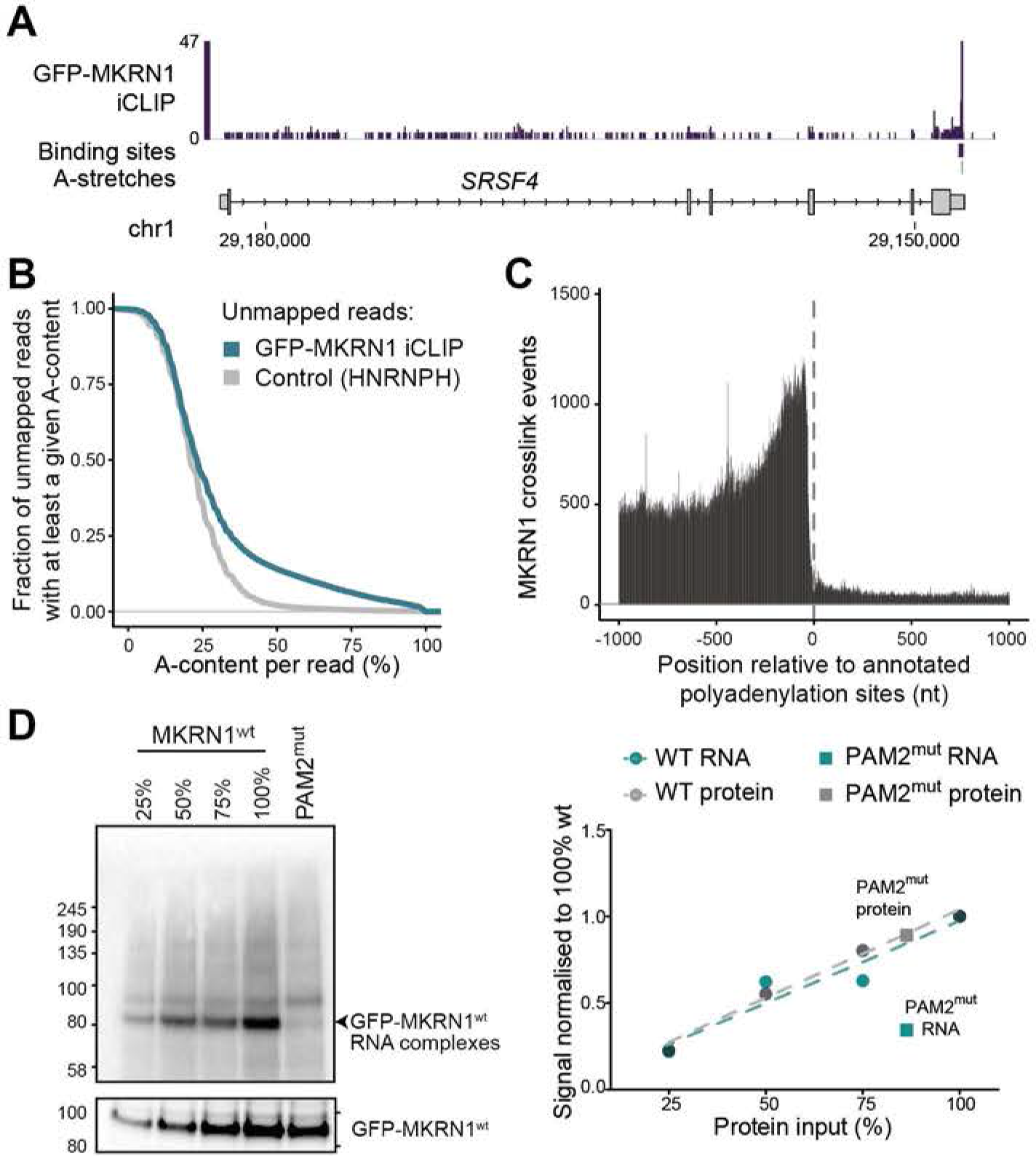
MKRN1 binds at poly(A) tails. (*A*) MKRN1 binds near the polyadenylation site of the *SRSF4* gene. Genome browser view as in **Fig. 2E**. (*B*) Unmapped MKRN1 iCLIP reads display increased A-content (more than half of all nucleotides in the read), evidencing poly(A) tail binding. Cumulative fraction of iCLIP reads (y-axis, merged replicates) that could not be mapped to the human genome (see Materials and methods) and show at least a given A-content (x-axis). iCLIP data for the unrelated RBP HNRNPH (Braun et al. 2018) are shown for comparison. (*C*) MKRN1 crosslink events increase towards 3’UTR ends. Metaprofile shows the sum of crosslink events per nt in a 2001-nt window around annotated polyadenylation sites of transcripts with >1 kb 3’ UTRs (n = 11,257). (*D*) Overall RNA binding of MKRN1 is strongly reduced when abrogating PABP interaction. Audioradiograph (left) of UV crosslinking experiments (replicate 1, with 4SU and UV crosslinking at 365 nm; replicates 2 and 3 in **Supplemental Fig. S5**) comparing GFP-MKRN1^PAM2mut^ with GFP-MKRN1^wt^ at different dilution steps for calibration. Quantification of radioactive signal of protein-RNA complexes and corresponding Western blots shown on the right. Uncropped gel images are shown in **Supplemental Fig. S10**.

### MKRN1 promotes ribosome stalling at poly(A) sequences

As outlined above, our iCLIP data evidenced that MKRN1 marks the beginning of poly(A) tails. Hence, it is conceivable that MKRN1 will also bind upstream of premature polyadenylation events within open reading frames. Based on MKRN1’s binding pattern, its interaction partners and its previously reported association with ribosomes (Simsek et al. 2017), we hypothesised that MKRN1 may be involved in the clearance of such transcripts by ribosome-associated quality control (RQC). In this process, ribosomes that translate into a poly(A) sequence, for instance upon stop codon readthrough and premature polyadenylation, are stalled and eventually recycled (Brandman and Hegde 2016; Joazeiro 2017). To test this hypothesis, we employed a recently introduced flow cytometry-based assay that monitors ribosome stalling in a dual fluorescence reporter (Juszkiewicz and Hegde 2017) (**Fig. 4A**).

**Figure 4.**
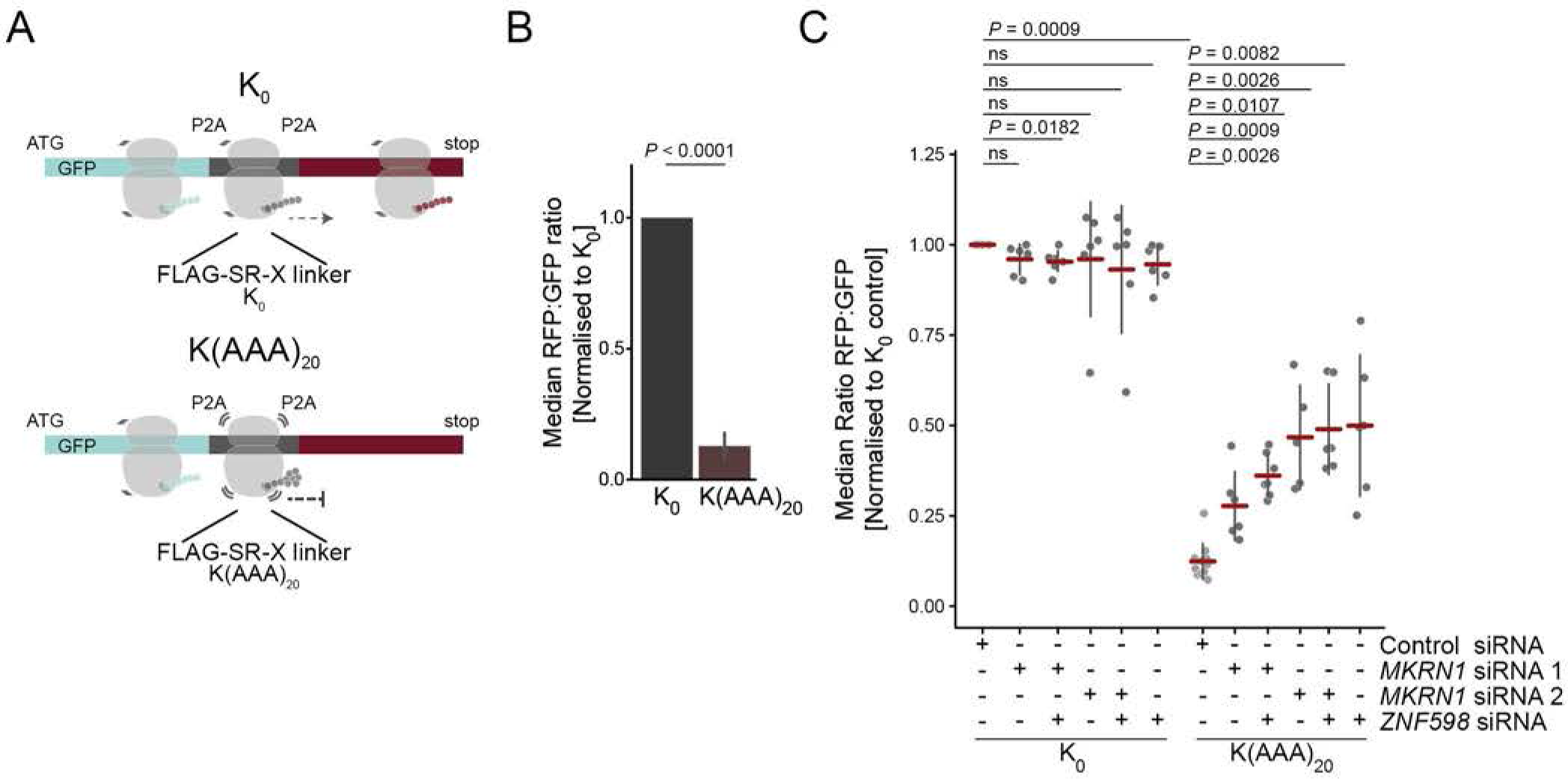
MKRN1 stalls ribosomes at poly(A) sequences. (*A*) The dual fluorescence reporter harbours an N-terminal GFP, followed by a FLAG-SR-X linker and a C-terminal RFP, which are separated by P2A sites to ensure translation into three separate proteins (Juszkiewicz and Hegde 2017). The resulting GFP:RFP ratio was determined using flow cytometry. The inserted fragment K(AAA)_20_ encodes 20 lysines by repeating the codon AAA. The starting vector without insert (K_0_) served as control. Schematic ribosomes illustrate translation of the respective reporter segments. (*B*) Ribosomes are efficiently stalled at K(AAA)_20_ in HEK293T cells. Median RFP:GFP ratios, normalised to K_0_, are shown. Error bars represent standard deviation of the mean (s.d.m., n = 6 replicates). *P* value indicated above (paired two-tailed t-test). (*C*) Ribosomes fail to stall in the absence of MKRN1. HEK293T cells were transfected with control siRNA or siRNAs targeting *MKRN1* (KD1 and KD2) or *ZNF598* for 24 h, followed by transfection of the reporter plasmids for 48 h. Western blots for KDs are shown in **Supplemental Fig. S6B**. RFP and GFP signals were analysed by flow cytometry. Median RFP:GFP ratios, normalised to K_0_ in control, are shown. Error bars represent s.d.m.; *P* values indicated above (paired two-tailed t-test, Benjamini-Hochberg correction, n ≥ 6 replicates; ns, not significant).

As reported previously, inserting a K(AAA)20 linker (encoding for 20 lysine residues) into the reporter resulted in predominant ribosome stalling compared to the starting vector (K_0_, **Fig. 4B** and **Supplemental Fig. 6A**). Importantly, *MKRN1* depletion with two independent siRNA sequences led to a reproducible recovery of RFP expression downstream of K(AAA)_20_, demonstrating that many ribosomes failed to stall at K(AAA)_20_ (*MKRN1* KD1 and KD2; **Fig. 4C** and **Supplemental Fig. S6A,B**). *MKRN1* KD2 seemed slightly more effective, possibly because this siRNA simultaneously decreased the transcript levels of the close paralogue *MKRN2* (**Supplemental Fig. S6C**). Notably, *MKRN1* KD2 impaired ribosome stalling to a similar level as KD of *ZNF598*, the E3 ubiquitin ligase that was recently reported to function in RQC (Garzia et al. 2017; Juszkiewicz and Hegde 2017; Sundaramoorthy et al. 2017). Moreover, simultaneous depletion of *MKRN1* and *ZNF598* was not additive, indicating that both proteins are necessary for function (**Fig. 4C** and **Supplemental Fig. S6A**). In addition, we noted a certain level of cross-regulation, such that *ZNF598* expression was decreased in *MKRN1* KD1 (but not in *MKRN1* KD2), whereas *ZNF598* overexpression reduced *MKRN1* expression (**Supplemental Fig. S6D,E**). Taken together, we conclude that MKRN1 contributes to efficient ribosome stalling in RQC.

### MKRN1 mediates the ubiquitylation of ribosome-associated proteins

RQC builds on a series of ubiquitylation events by multiple E3 ubiquitin ligases, including Listerin and ZNF598 (Brandman and Hegde 2016). In order to identify putative ubiquitylation substrates of MKRN1, we first determined the protein interactome of the ligase-deficient mutant GFP-MKRN1^RINGmut^. In three replicate experiments, we quantified 1,097 protein groups present in at least two out of three replicates (**Supplemental Table S1**), revealing 137 proteins that were significantly enriched compared to GFP-MKRN1^wt^ (**Supplemental Fig. S7**). Intriguingly, these included RPS10, a ribosomal protein that was previously reported to be modified by ZNF598 during RQC (Garzia et al. 2017; Juszkiewicz and Hegde 2017; Sundaramoorthy et al. 2017).

In order to directly test for ubiquitylation of putative substrates of MKRN1, we performed ubiquitin remnant profiling to compare the relative abundance of di-glycine-modified lysines in wild type and *MKRN1* KD cells. We quantified 2,324 ubiquitylation sites (in 1,264 proteins) that were detected in all four replicate experiments (**Supplemental Table S3**). Notably, *MKRN1* depletion led to a significantly decreased abundance of 29 ubiquitylation sites on 21 proteins (FDR < 10%, **Fig. 5A**). The majority of the ubiquitylation targets assembled into a coherent cluster of translational regulators based on previously reported protein-protein interactions and functional annotations (**Fig. 5B,C** and **Supplemental Fig. 8A**). Among these proteins, we had already detected PABPC1/4, IGF2BP1, ELAVL1, MOV10, LARP1, and RPS10 as significant interactors of GFP-MKRN1^wt^ and/or GFP-MKRN1^RINGmut^ (**Fig. 1A**, **Fig. 5F** and **Supplemental Fig. S7**). Importantly, we detected a significant decrease in ubiquitylation at lysine 107 of RPS10 (K107; **Fig. 5D**). In order to distinguish differential ubiquitylation from protein level changes, we also measured the total protein levels in *MKRN1* KD cells and did not observe changes in RPS10, PABPC1/4, IGF2BP1/2/3, ELAVL1, and MOV10 protein levels (**Supplemental Fig. 8B** and **Supplemental Table S4**). Taken together, we conclude that MKRN1 mediates ubiquitylation of the ribosomal protein RPS10 and several translational regulators during ribosome-associated quality control.

**Figure 5.**
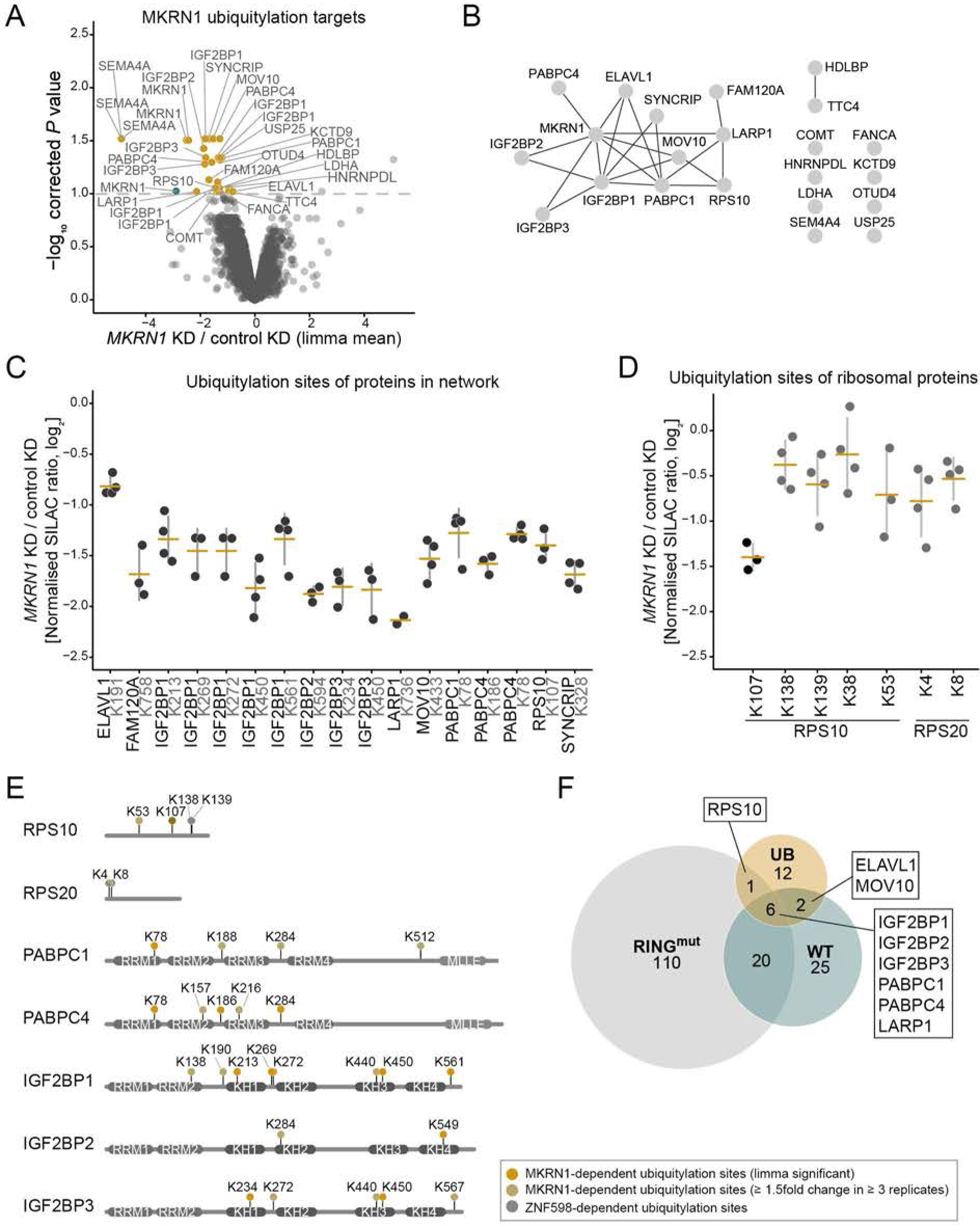
MKRN1 ubiquitylates ribosomal protein RPS10 and translational regulators. Ubiquitin remnant profiling to compare the relative abundance of ubiquitylation sites in *MKRN1* KD2 and control HEK293T cells. Ubiquitin remnant peptides were enriched and analysed by quantitative mass spectrometry, quantifying a total of 15,528 ubiquitylation sites on 4,790 proteins. 29 putative MKRN1 target sites with significantly decreased ubiquitylation upon *MKRN1* KD2 (FDR < 10%, n = 4 replicates) are highlighted and labelled with the respective protein name. Note that many proteins contain several differentially regulated ubiquitylation sites. (*B*) Protein interaction network of 21 proteins with putative MKRN1 ubiquitylation target sites (significantly reduced, shown in (*A*)). The functional interactions were obtained from the STRING and BioGrid databases and our study. Visualisation by Cytoscape. (*C*) Ubiquitin remnant profiling results for significantly regulated ubiquitylation sites (FDR < 10%) in proteins from network in (*B*). Mean and standard deviation of the mean (s.d.m., error bars) are given together with all data points. (*D*) Ubiquitin remnant profiling results for seven quantified ubiquitylation sites in RPS10 and RPS20. Significant changes are shown in black (FDR < 10%) and non-significant changes in grey. Representation as in (*C*). (*E*) Comparison of ubiquitylation sites in selected target proteins that are modified by ZNF598 and MKRN1 during RQC. (*F*) Comparison of enriched proteins from the interactomes for GFP-MKRN1^wt^ (over GFP, see **Fig. 1A**) and GFP-MKRN1^RINGmut^ (over GFP-MKRN1^wt^, see **Supplemental Fig. S7B**) with the proteins containing MKRN1 ubiquitylation targets sites (UB, see (*A*)). Protein names of overlapping targets are given.

## Discussion

Ribosome-associated quality control is essential to recognise and clear terminally stalled ribosomes. Here, we uncover MKRN1 as a novel factor in RQC. Our data indicate that MKRN1 is positioned upstream of poly(A) sequences through direct interaction with PABP, thereby marking the beginning of poly(A) tails. We propose that in case of premature polyadenylation, MKRN1 stalls the translating ribosome and initiates RQC by ubiquitylating ribosomal protein RPS10, PABP and other translational regulators (**Fig. 6**).

**Figure 6.**
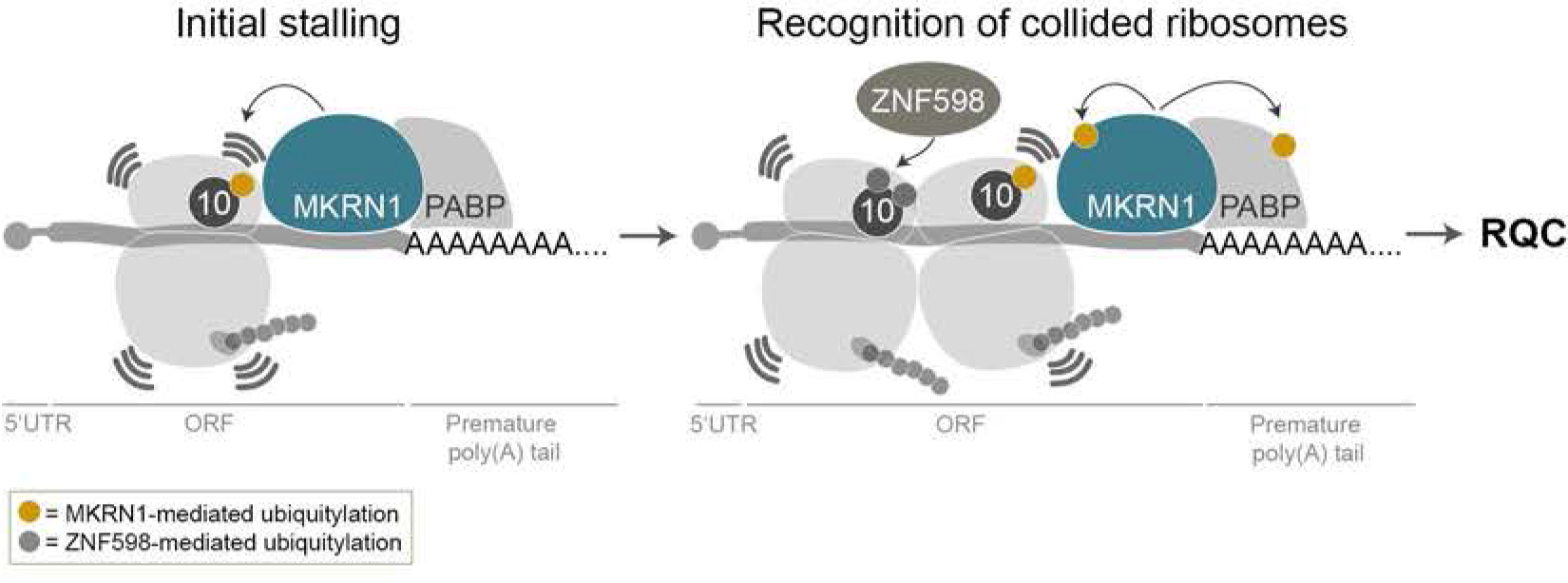
MKRN1 is a sensor for poly(A) sequences that stalls ribosomes to initiate ribosome-associated quality control. Proposed model of MKRN1 function: MKRN1 is positioned upstream of (premature) poly(A) tails via interaction with PABP. Ribosomes translating the open reading frame run into MKRN1 that acts as a roadblock to prohibit poly(A) translation. Upon contact with the translating ribosome, MKRN1 ubiquitylates the 40S ribosomal protein RPS10. This stalls the ribosome, causing the trailing ribosomes to collide. ZNF598 recognises the collided ribosomes and ubiquitylates ribosomal proteins to promote RQC.

### PABP recruits MKRN1 upstream of A-stretches and poly(A) tails

Central to our model is the specific RNA-binding behaviour of MKRN1, which is recruited to mRNA by PABP to mark the beginning of poly(A) tails. This builds on the following observations: (i) We and others show that MKRN1 and PABP interact via the PAM2 motif (Miroci et al. 2012). (ii) MKRN1 binding to RNA is strongly reduced when interaction with PABP is abolished. (iii) The association of strong MKRN1 binding with continuous A-runs of ≥8 A’s mirrors the footprint of one RNA recognition motif (RRM) domain of PABP, indicating that the binding of one RRM to poly(A) is sufficient for MKRN1 recruitment (Webster et al. 2018). On such short A-stretches, MKRN1 might stabilise PABP binding, while on longer A-stretches, PABP might be the major driving force to recruit MKRN1. This interaction might also anchor the first PABP at the beginning of the poly(A) tail. One possible function could be the stabilisation of PABP on short poly(A) tails to promote efficient translation (Lima et al. 2017). In yeast, where a MKRN1 ortholog is missing (see below), this anchoring is thought to be achieved by Pab1p itself via its fourth RRM domain (Webster et al. 2018). Of note, a parallel study with the Mkrn1 ortholog from *D. melanogaster* demonstrates binding of a Mkrn1/pAbp complex at an A-stretch in the 3’ UTR of *oskar* mRNA, which is involved in translational control and required for oogenesis (Dold et al., parallel submission; preprint available at bioRxiv, doi: 10.1101/501643).

### MKRN1 ubiquitylates RPS10 and translational regulators to stall ribosomes

Our data suggest that ribosomes encountering the MKRN1-PABP complex are stalled, possibly via ubiquitylation of RPS10 and other MKRN1 interactors. Concordantly, ZNF598, a factor that was recently shown to function in RQC, was also found to mediate ubiquitylation of RPS10 (Juszkiewicz et al. 2018). In conjunction with its unique RNA-binding behaviour, we therefore hypothesise that MKRN1 acts as a first line of defence against poly(A) translation. We propose that MKRN1 is recruited by PABP to the beginning of poly(A) tails, including premature polyadenylation events within open reading frames, where it represents a physical “roadblock” to the translating ribosome. Upon contact with the translating ribosome, MKRN1 ubiquitylates K107 on RPS10, thereby stalling the ribosome before it translates the poly(A) tail. Subsequently, the trailing ribosomes collide with the initially stalled ribosome. ZNF598 recognises the collision interface and ubiquitylates the collided ribosomes (Simms et al. 2017; Juszkiewicz et al. 2018). In summary, we suggest that a sequence of MKRN1-mediated and ZNF598-mediated ubiquitylation events on ribosomal proteins and possibly other factors, including PABPC1, triggers ribosome-associated quality control.

### Differences between human and yeast RQC explain the requirement for MKRN1

Many known components of the RQC machinery, such as Listerin (Ltn1p in yeast) and ZNF598 (Hel2p in yeast), are identical from yeast to human, however the molecular signals that are recognised differ partially. In yeast, RQC can be triggered by an excess of positively charged amino acids (lysine and arginine), which are sensed while they pass through the ribosomal exit tunnel (Lu and Deutsch 2008; Letzring et al. 2013). In contrast, in human, sensing the aberrant mRNAs does not occur via the encoded amino acids but at the level of the mRNA sequence and corresponding tRNAs, such that only poly(A) effectively results in ribosome stalling (Arthur et al. 2015; Garzia et al. 2017; Juszkiewicz and Hegde 2017). We propose that MKRN1 acts as direct reader of poly(A) sequences based on its interaction with PABP. Consistent with this conceptual difference, there is no functionally equivalent ortholog of MKRN1 in yeast (Yth1p and Lee1p are similar, but lack RING domain and PAM2 motif; **Supplemental Fig. 1C**). Why yeast and human employ partially different mechanisms to detect poly(A) translation is currently unclear, but it has been suggested that spurious translation of poly-lysine stretches from long human poly(A) tails might target the aberrant proteins to the nucleus (Juszkiewicz and Hegde 2017). Loss of mRNA surveillance and RQC deficiency can lead to protein aggregation and culminate in proteotoxic stress, which in turn is lined to neurological disorders such as amyotrophic lateral sclerosis (Choe et al. 2016; Jamar et al. 2018). Hence, recognition of poly(A) sequences prior to their translation might be particularly beneficial in humans.

## Materials and methods

### Cell culture

HEK293T cells were obtained from DSMZ and cultured in DMEM (Life Technologies) with 10% fetal bovine serum (Life Technologies), 1% penicillin/streptomycin (Life Technologies), and 1% L-glutamine (Life Technologies). All cells were maintained at 37°C in a humidified incubator containing 5% CO_2_ and routinely tested for mycoplasma infection. For SILAC labelling, cells were maintained in media containing either L-arginine and L-lysine (light SILAC label), L-arginine (^13^C_6_) and L-lysine (^2^H_4_) (medium SILAC label), or L-arginine (^13^C_6_-^15^N_4_) and L-lysine (^13^C_6_-^15^N_2_) (heavy SILAC label) (Cambridge Isotope Laboratories).

### Vectors

The following vectors, suitable for Gateway Cloning, were obtained either from the IMB Core Facility ORFeome Collection (Collaboration 2016) or from the Harvard PlasmID Repository (https://plasmid.med.harvard.edu/PLASMID/): pENTR221-MKRN1, pENTR221-PABPC1, pENTR223.1-IGF2BP1, pENTR221-ELAVL1, pCMV-SPORT-ZNF598. Coding sequences from the entry vectors were cloned into the mammalian expression vectors pMX-DEST53-IP-GFP by LR Gateway cloning according to the manufacturer’s recommendations (Gateway LR Clonase II Enzyme mix; Life Technologies). Dual fluorescence reporter plasmids (pmGFP-P2A-K_0_-P2A-RFP, pmGFP-P2A-(K^AAA^)_12_-P2A-RFP, pmGFP-P2A-(K^AAA^)_20_-P2A-RFP, and pmGFP-P2A-(R^CGA^)_10_-P2A-RFP) were generously provided by Ramanujan S. Hegde (MRC Laboratory of Molecular Biology, Cambridge, UK) (Juszkiewicz and Hegde 2017).

### Cloning

All MKRN1 mutant plasmids were generated with the Q5 Site-Directed Mutagenesis Kit (NEB) according to the manufacturer’s recommendations. In order to disrupt MKRN1’s interaction with PABP (MKRN1^PAM2mut^), three point mutations were introduced into the PAM2 motif (A169S, F172A, P174A; **Fig. 1B**) as previously described (Pohlmann et al. 2015). In MKRN1^RINGmut^, a previously described mutation in the RING domain (H307E) was introduced to abolish E3 ubiquitin ligase function (Kim et al. 2005). All primers used for introducing mutations into MKRN1 are listed in **Supplemental Table S5**.

### Transfections

Overexpression of vectors was performed using Polyethylenimine MAX 4000 (Polysciences, 24885-2) with a DNA:PEI ratio of 1:10. Knockdowns were performed with siRNAs (**Supplemental Table S6**) using Lipofectamine RNAiMAX (Life Technologies) according to the manufacturer’s recommendations.

### Affinity purification (AP) for Western blot analyses

GFP-based affinity purifications (APs) were performed as described before (Hildebrandt et al. 2017). In brief, HEK293T cells transiently expressing GFP (empty vector) or a GFP-tagged target protein were used. The cells were lysed in modified RIPA (mRIPA) buffer supplemented with protease inhibitors (protease inhibitor cocktail, Sigma), 1 mM sodium orthovanadate, 5 mM β-glycerophosphate, 5 mM sodium fluoride, and 10 mM N-ethylmaleimide (NEM) (all from Sigma). Protein concentrations were determined using the Pierce BCA Protein Assay Kit (Thermo Fisher). GFP-trap agarose beads (Chromotek) were incubated with the cleared lysate for 1 h at 4°C. After five washes with mRIPA buffer, the beads were resuspended in LDS sample buffer (Life Technologies) and heated to 70°C for 10 min. For RNase digests, the enriched proteins were incubated with 0.5 U/μl RNase A (Qiagen) and 20 U/μl RNase T1 (Thermo Fisher Scientific) for 30 min at 4°C after the first two washes in mRIPA buffer.

### Sample preparation for the protein interactome analysis

GFP-based APs were performed as described before (Hildebrandt et al. 2017). In brief, HEK293T cells transiently expressing GFP (empty vector) were cultured in light SILAC medium, while cells expressing N-terminally GFP-tagged MKRN1 wt or mutants were cultured in medium or heavy SILAC medium. The cells were lysed as described above. After washing in mRIPA buffer, GFP-trap agarose beads were incubated with the cleared lysate for 1 h at 4°C. All AP samples were washed four times with mRIPA buffer, combined and washed again in mRIPA buffer. The beads were heated in LDS sample buffer, supplemented with 1 mM dithiothreitol (DTT; Sigma, D5545) for 10 min at 70°C and alkylated using 5.5 mM 2-chloroacetamide (CAA; Sigma, C0267) for 30 min at RT in the dark (Nielsen et al. 2008).

### Sample preparation for the proteome analysis

*MKRN1* KD using siRNA2 was performed in heavy labelled SILAC cells and control KD was performed in light labelled SILAC cells in two replicates. For the third replicate, a label swop was performed, knocking down *MKRN1* (siRNA2) in light labelled SILAC cells and control in heavy labelled SILAC cells. For proteome analysis, cells were lysed as described above. Subsequently, 25 μg protein from each SILAC condition (50 μg in total) were pooled and processed as described below.

### Sample preparation for mass spectrometry

The enriched proteins were resolved by SDS-PAGE on a NuPAGE 4-12% Bis-Tris protein gel (Thermo Fisher Scientific) and stained using the Colloidal Blue Staining Kit (Life Technologies). Proteins were in-gel digested using trypsin, before peptides were extracted from the gel. To concentrate, clear and acidify the peptides, they were bound to C18 StageTips as described previously (Rappsilber et al. 2007).

### Mass spectrometry data acquisition

Peptide fractions were analysed on a quadrupole Orbitrap mass spectrometer (Thermo Q Exactive Plus, Thermo Scientific) coupled to an uHPLC system (EASY-nLC 1000, Thermo Scientific) (Michalski et al. 2011). Peptide samples were separated on a C18 reversed phase column (length: 20 cm, inner diameter: 75 μm, bead size: 1.9 μm) and eluted in a linear gradient from 8 to 40% acetonitrile containing 0.1% formic acid in 105 min for the interactome analyses, in 175 min for the proteome analyses, or in 125 min for the ubiquitylome analyses. The mass spectrometer was operated in data-dependent positive mode, automatically switching between MS and MS^2^ acquisition. The full scan MS spectra (m/z 300–1650) were acquired in the Orbitrap. Sequential isolation and fragmentation of the ten most abundant ions was performed by higher-energy collisional dissociation (HCD) (Olsen et al. 2007). Peptides with unassigned charge states, as well as with charge states less than +2 were excluded from fragmentation. The Orbitrap mass analyser was used for acquisition of fragment spectra.

### Peptide identification and quantification

Raw data files were analysed and peptides were identified using the MaxQuant software (version 1.5.28) (Cox et al. 2009). Parent ion and MS^2^ spectra were compared to a database containing 92,578 human protein sequences obtained from UniProtKB (release June 2018), coupled to the Andromeda search engine (Cox et al. 2011). Cysteine carbamidomethylation was set as a fixed modification. N-terminal acetylation, oxidation, and N-ethylmaleimide (NEM) were set as variable modifications. For ubiquitylome data analysis, glycine-glycine (GlyGly) modification of lysine was additionally set as a variable modification. The mass tolerance for the spectra search was set to be lower than 6 ppm in MS and 20 ppm in HCD MS^2^ mode. Spectra were searched with strict trypsin specificity and allowing for up to three mis-cleavages. Site localisation probabilities were determined by MaxQuant using the PTM scoring algorithm as described previously (Elias and Gygi 2007; Cox and Mann 2008). Filtering of the dataset was based on the posterior error probability to arrive at a false discovery rate (FDR) < 1% estimated using a target-decoy approach. Proteins that were categorised as “only identified by site”, potential contaminants and reverse hits were removed. Only proteins identified with at least two peptides (including at least one unique peptide) and a SILAC ratio count of at least two were used for analysis. For AP experiments, proteins that were quantified in at least two out of three experiments were kept for further analysis. In total, we quantified 1,106 and 1,097 protein groups in the AP experiments with GFP-MKRN1^wt^ (**Fig. 1A**), GFP-MKRN1^PAM2mut^ (**Fig. 1D**) and GFP-MKRN1^RINGmut^ (**Supplemental Fig. S7**), respectively (**Supplemental Table S1**). The SILAC ratios were log_2_ transformed and converted into an asymmetric z-score based on the mean and interquartile range of the distribution as described previously (Cox and Mann 2008). For statistical analysis, a moderated t-test from the limma algorithm was used (Ritchie et al. 2015). Enriched proteins with an FDR < 5% were determined to be significantly enriched interactors (for GFP-MKRN1^wt^). For proteins enriched in GFP-MKRN1^RINGmut^ over GFP-MKRN1^wt^, proteins with an FDR < 5% and a GFP-MKRN1^wt^/GFP z-score > 1 were selected. In the proteome experiment, we quantified 6,439 protein groups, present in all three replicates. Ratio-ratio and ratio-intensity plots were created in R (version 3.4.3) using RStudio (http://www.rstudio.com/).

### Functional annotation of MKRN1 interactors and MKRN1-ubiquitylation targets

In order to assess the functions of MKRN1-interacting proteins and proteins with MKRN1-dependent ubiquitylation sites, we performed gene ontology (GO) enrichment analyses using the Database for Annotation, Visualization and Integrated Discovery (DAVID 6.7) for three GO domains (Jiao et al. 2012). Enriched GO terms (modified Fisher exact test, adjusted *P* value < 0.05, Benjamini-Hochberg correction; **Supplemental Fig. S2B,** **S8A**) were visualised using REVIGO (Reduce & Visualize Gene Ontology) allowing medium GO term similarity (Supek et al. 2011).

### Western blot

Denatured proteins were separated by SDS-PAGE on a NuPAGE 4-12% Bis-Tris protein gel (Life Technologies) and transferred to a 0.45 μm nitrocellulose membrane (VWR). For detection, either fluorophore-coupled secondary antibodies or HRP-conjugated secondary antibodies and WesternBright Chemiluminescent Substrate (Biozym Scientific) or SuperSignal West Pico Chemiluminescent Substrate (Life Technologies) were used. Western blots were quantified by determining the background-subtracted densities of the protein of interest using ImageJ (Schindelin et al. 2015). The signal from the AP (against GFP-tagged protein of interest) was normalised to the respective control samples expressing the empty vector or to the input.

### Antibodies

The following antibodies were used: anti-GFP (B-2 clone; Santa Cruz; sc-9996), anti-MKRN1 (Bethyl Laboratories, A300-990A), anti-PABPC1/3 (Cell Signaling, 4992), anti-Znf598 (N1N3; GeneTex; GTX119245), anti-αTubulin (Sigma Aldrich, T-5168), anti-Rabbit IgG (Cell Signaling; 7074), anti-Mouse IgG (Cell Signaling; 7076), IRDye^®^ 680RD Goat anti-Mouse IgG (P/N 925-68070), and IRDye^®^ 800CW Goat anti-Rabbit IgG (P/N 925-32211) (both LI-COR Biosciences GmbH).

### RNA isolation, cDNA synthesis and qPCR

Cells were washed twice in ice-cold PBS and harvested. RNA was isolated using the RNeasy Plus Mini Kit (Qiagen) according to the manufacturer’s recommendations. 500 ng total RNA was transcribed into cDNA using random hexamer primers (Thermo Scientific) and the RevertAid Reverse Transcriptase (Thermo Scientific) according to the manufacturer’s recommendations. qPCR was performed using the Luminaris HiGreen qPCR Master Mix, low ROX (Thermo Scientific) according to the manufacturer’s recommendations with 10 μM forward and reverse primers (**Supplemental Table S5**).

### iCLIP experiments and data processing

iCLIP libraries were prepared as described previously (Huppertz et al. 2014; Sutandy et al. 2016). HEK293T cells ectopically expressing either GFP alone (empty vector) or N-terminally GFP-tagged MKRN1 wild type (GFP-MKRN1^wt^), GFP-MKRN1^PAM2mut^, or GFP-MKRN1^RINGmut^ were used. For crosslinking, confluent cells were irradiated once with 150 mJ/cm^2^ at 254 nm in a Stratalinker 2400 or treated with 4-thiouridine (100 μM for 16 h) and irradiated with 3x 300 mJ/cm^2^ in a Stratalinker 2400 with 365 nm bulbs. For IP, 10.5 μg anti-GFP antibody (goat, Protein Unit, MPI-CBG, Dresden) were used per sample. The libraries were sequenced as 50-nt single-end reads on an Illumina MiSeq platform (**Supplemental Table S2**).

Basic sequencing quality checks were applied to all reads using FastQC (version 0.11.5) (https://www.bioinformatics.babraham.ac.uk/projects/fastqc/). Afterwards, reads were filtered based on sequencing qualities (Phred score) of the barcode region. Only reads with at most one position with a sequencing quality < 20 in the experimental barcode (positions 4 to 7) and without any position with a sequencing quality < 17 in the random barcode (positions 1-3 and 8-9) were kept for further analysis. Remaining reads were de-multiplexed based on the experimental barcode on positions 4 to 7 using Flexbar (version 3.0.0) (Dodt et al. 2012) without allowing mismatches.

All following steps of the analysis were performed on all individual samples after de-multiplexing. Remaining adapter sequences were trimmed from the right end of the reads using Flexbar (version 3.0.0) allowing up to one mismatch in 10 nt, requiring a minimal overlap of 1 nt of read and adapter. After trimming off the adapter, the barcode is trimmed off of the left end of the reads (first 9 nt) and added to the header of the read, such that the information is kept available for downstream analysis. Reads shorter than 15 nt were removed from further analysis.

Trimmed and filtered reads were mapped to the human genome (assembly version GRCh38) and its annotation based on GENCODE release 25 (Harrow et al. 2012) using STAR (version 2.5.4b) (Dobin et al. 2013). When running STAR, up to two mismatches were allowed, soft-clipping was prohibited and only uniquely mapping reads were kept for further analysis.

Following mapping, duplicate reads were marked using the dedup function of bamUtil (version 1.0.13), which defines duplicates as reads whose 5’ ends map to the same position in the genome (https://github.com/statgen/bamUtil). Subsequently, marked duplicates with identical random barcodes were removed since they are considered technical duplicates, while biological duplicates showing unequal random barcodes were kept.

Resulting bam files were sorted and indexed using SAMtools (version 1.5) (Li et al. 2009). Based on the bam files, bedgraph files were created using bamToBed of the BEDTools suite (version 2.25.0) (Quinlan and Hall 2010), considering only the position upstream of the 5' mapping position of the read, since this nucleotide is considered as the crosslinked nucleotide. bedgraph files were then transformed to bigWig file format using bedGraphToBigWig of the UCSC tool suite (Kent et al. 2010).

### Identification and characterisation of MKRN1 binding sites

Peak calling was performed on merged iCLIP coverage tracks (crosslink events per nucleotide) from the three replicates based on GENCODE annotation (release 27, GRCh38) using ASPeak (version 2.0; default setting plus ‒nornaseq to estimate parameters p and r for the negative binomial distributions in a 500-nt window around each peak) (Kucukural et al. 2013). The initially predicted peaks were resized to uniform 9-nt windows around their weighted centred as defined by ASPeak. To avoid artefacts, we removed sparsely covered peaks that harbour crosslink events on less than three nucleotides within the 9-nt region window. We iteratively merged all remaining windows if overlapping by at least 1 nt, by defining the position with the cumulative half maximum count of crosslink events as new window centre. We further kept only reproducible windows with at least three crosslink events from any two replicates. Finally, we excluded all windows overlapping with none or multiple protein-coding genes (GENCODE annotations support level ≥ 2 and transcript support level ≥ 3), and assign each binding site to a distinct genomic region (3’ UTR, 5’ UTR, CDS, intron). Consistent with the mostly cytoplasmic localisation of MKRN1 (Miroci et al. 2012; Cassar et al. 2015; Hildebrandt et al. 2017), less than 6% of the binding sites were predicted within introns, which were excluded from further analysis. This procedure yielded a total of 7,331 MKRN1 binding sites in 2,163 genes.

In order to estimate binding site strength and to facilitate comparisons between binding sites (**Fig. 2B,D** and **Supplemental Fig. 3D-F**, **4A,C,D**), we corrected for transcript abundance by representing the crosslink events within a binding site as a ‘signal-over-background’ ratio (SOB). The respective background was calculated as the sum Of crosslink events outside of binding sites (plus 5 nt to either side) by the merged length of all exons. 3’ UTR lengths were restricted to 10 nt past the last MKRN1 binding site or 500 nt if no binding site was present. SOB calculations were performed separately for each replicate and then averaged. No SOB value was assigned for genes with a background of < 10 crosslink events, resulting in SOB values for 97% of all binding sites.

In order to assess the local RNA sequence context of MKRN1 binding sites (**Fig. 2B** and **Supplemental Fig. S4A**), enriched 4-mers were counted inside the 9-nt binding sites as well as within 40-nt before and after. To estimate an empirical background distribution, 1,000 9-nt windows were randomly picked in 3’ UTRs and 4-mer frequencies were counted in the same windows. This process was repeated 100 times, and the resulting mean and standard deviation were used to calculate the z-score for each 4-mer.

In order to define the A-rich regions downstream of MKRN1 binding sites in 3’ UTRs (A-rich stretches), we used a maximisation approach in a 55-nt search space starting from the binding site centre. Within this space, we calculated the percentage of A nucleotides (A-content) for windows of increasing size (8-30 nt) and selected the stretch with highest value for each window size. In case of ties, the window closer to the binding site was preferred, resulting in a set of 23 candidate A-stretches with the maximal A-content for each length. Next, we computed the longest continuous A run (LCA) and a weighted A-content (multiplying the A-content with the number of A nucleotides) for each candidate A-stretch. Candidate A-stretches with an A-content < 70%, a weighted A-content < 11 and an LCA < 4 were excluded. The final A-stretch for each binding site was then selected in a hierarchical manner, preferring LCA over weighted A-content. Lastly, overlapping A-stretches of neighbouring binding sites were merged by selecting the highest scoring A-stretch, based on LCA and weighted A-content. In total, this procedure identified 1,412 non-overlapping A-stretches, associated with 1,848 binding sites.

In order to estimate the extent of MKRN1 binding to poly(A) tails (**Fig. 3B**), we evaluated the percentage of adenosine within the iCLIP reads that could not be mapped to the human genome without soft-clipping (see above). iCLIP data for heterogeneous nuclear ribonucleoprotein H (HNRNPH) served as control (Braun et al. 2018). Annotated transcript 3’ ends (i.e. polyadenylation sites) were taken from GENCODE (all annotated protein-coding transcripts with support level ≤ 2 and transcript support level ≤ 3; release 28, GRCh38.p12; www.gencodegenes.org). For **Fig. 3C**, all crosslink events within a 2-kb window around the polyadenylation sites for 3’ UTR longer than 1 kb were counted.

### Evolutionary characterisation of Makorin protein family

Four different ortholog searches were performed using HaMStR-OneSeq (Ebersberger et al. 2014) against the Quest for Orthologs Consortium protein set, containing 78 species (release 2017_04) (Sonnhammer et al. 2014). For each run, a different seed protein was chosen: human MKRN1-3 (UniProt identifiers Q9UHC7, Q9H000 and Q13064) and MKRN4 from zebrafish (A9C4A6). In order to identify proteins with a similar domain architecture, we calculated a unidirectional feature architecture similarity (FAS) score which compares the domain architecture of the seed protein and the predicted ortholog (Koestler et al. 2010). Predicted orthologues with FAS < 0.7 were removed after initial assessment. Finally, all vertebrate species and selected invertebrate species were used for reconstruction of a maximum likelihood (ML) tree. For this, protein sequences were aligned using MAFFT v7.294b L-INS-i (Katoh and Standley 2013), and ML trees with 100 bootstrap replicates were calculated using RAxML version 8.1.9 (Stamatakis 2014). Settings for a rapid bootstrap analysis and searching for the best scoring ML tree in one program run (-f a) and an automatic selection of the best fitting amino acid substitution model (-m PROTGAMMAAUTO) were chosen. Reconstructed trees were visualised using FigTree v1.4.2 (http://tree.bio.ed.ac.uk/software/figtree/).

The phylogenetic tree and FASTA sequences from the ortholog dataset were loaded into DoMosaics (http://www.domosaics.net) and Pfam domains were annotated with HMMER (http://hmmer.org/, default parameters). Since the PAM2 motif in all Makorin proteins differs from the described consensus motif (Albrecht and Lengauer 2004), a custom Hidden Markov Model was trained on PAM2 motifs from selected Makorin orthologs and used for a HMMER scan of the orthologs (no E-value cutoff). The same procedure was repeated for the PAM2-like motif (PAM2L) (Pohlmann et al. 2015).

### Dual fluorescence translation stall assay via flow cytometry

Knockdowns were performed for 24 h, before the dual fluorescence reporter plasmids were ectopically expressed for 48 h. Cells were washed in PBS and trypsinised. After sedimentation, cells were resuspended in DPBS supplemented with 2 mM EDTA. Cellular GFP and RFP fluorescence was measured using flow cytometry on a LSRFortessa SORP (BD Biosciences). Data analysis was done using FlowJo (v10) (FlowJo, LLC). For statistical testing, paired two-tailed Student’s t-tests with Benjamini-Hochberg correction were performed on n ≥ 6 replicates.

### Ubiquitin remnant profiling

Di-glycine remnant profiling was performed as described before (Wagner et al. 2011; Heidelberger et al. 2018). In four different experiments, isotope labels were assigned as follows: experiment 1, *MKRN1* KD1 (siRNA1), *MKRN1* KD2 (siRNA2) and control siRNA with light, medium and heavy SILAC labels, respectively; experiment 2, *MKRN1* KD2 (siRNA2) and control siRNA with heavy and light SILAC labels, respectively; experiment 3, *MKRN1* KD2 (siRNA2) and control siRNA with heavy and light SILAC labels, respectively; experiment 3, *MKRN1* KD2 (siRNA2) and control siRNA with light and heavy SILAC labels, respectively. Cells were treated with the proteasome inhibitors bortezomib (1 μM, 8h, replicate 1; Santa Cruz Biotechnology) or MG132 (10 μM, 2 h, replicates 2, 3, 4; Sigma). Proteins were precipitated in acetone. Proteins were digested with endoproteinase Lys-C (Wako Chemicals) and sequencing-grade modified trypsin (Sigma). To purify the peptides, reversed-phase Sep-Pak C18 cartridges (Waters) were used. Modified peptides were enriched using di-glycine-lysine antibody resin (Cell Signaling Technology). The enriched peptides were eluted with 0.15% trifluoroacetic acid in water, then fractionated using micro-column-based strong-cation exchange chromatography (SCX) (Weinert et al. 2013) before being desalted on reversed-phase C18 StageTips (Rappsilber et al. 2007). Samples were analysed by quantitative mass spectrometry and MaxQuant as described above. To identify significantly regulated ubiquitylation sites, the limma algorithm was applied (Ritchie et al. 2015). A *P* value < 0.1 after multiple testing correction was used as a cut-off to determine up- and downregulated ubiquitylation sites. Volcano and dot plots were created in R (version 3.4.3).

### Functional interaction network of MKRN1 ubiquitylation target proteins

The functional protein interaction network analysis was performed by integrating interaction data from the STRING database (score > 0.4), the BioGrid database and our own findings (Franceschini et al. 2013; Chatr-Aryamontri et al. 2017). Cytoscape (version 3.6.1) was used to visualise the protein interaction network (Saito et al. 2012).

### Availability of data and materials

The mass spectrometry proteomics data have been deposited to the ProteomeXchange Consortium (http://proteomecentral.proteomexchange.org) via the PRIDE partner repository with the dataset identifier PXD011772.

Raw and processed iCLIP data are available at GEO under the accession number GSE122869.

## Acknowledgements

We would like to thank all members of the Zarnack, König and Beli labs, as well as René Ketting, Nadine Wittkopp, and Miguel Almeida for fruitful discussions. The authors gratefully thank the Ramanujan S. Hegde for providing the dual fluorescence reporter plasmids. We thank Anja Freiwald for assistance with mass spectrometry analysis and Dr. Stefan Simm for assistance with evaluating PAM2 mutations. The support of the IMB Core Facilities Bioinformatics, Flow Cytometry, Genomics, and the use of its Illumina MiSeq, as well as the DFG-funded Mass Spectrometer Q Exactive Plus (INST 247/766-1 FUGG) are gratefully acknowledged. K.Z. was supported by the LOEWE program Ubiquitin Networks (Ub-Net) of the State of Hesse (Germany) and the SFB 902 of the German Research Foundation. P.B. is supported by the Emmy Noether Program (BE 5342/1-1), the SFB 1177 of the German Research Foundation and the Marie Curie Career Integration Grant from the European Commission (grant agreement number: 630763). The project was funded by the German Research Foundation (DFG) as part of SPP1925 to J.-Y.R. (RO 4681/4-1) and J.K. (KO 4566/3-1). Animal shapes in Supplemental Fig. S1A were obtained from PhyloPic and are used under the Creative Common Attribution-NonCommercial-ShareAlike 3.0 Unported license.

## Author contributions

A.H. performed iCLIP experiments, flow cytometry measurements of dual fluorescence reporters and most proteomics experiments. M.B. performed most bioinformatics analyses of MKRN1 iCLIP data. C.R. analysed MKRN1 binding at polyadenylation sites and poly(A) tails. A.B. and S.B. performed initial iCLIP data processing and analysis. A.H. and A.B. analysed the proteomics data. J.B.H. and A.V. contributed to replicate ubiquitin remnant profiling experiments and AP-Western blot experiments, respectively. H.H. performed replicate iCLIP and replicate AP-Western blot experiments. C.R. and I.E. contributed evolutionary characterisation of Makorin proteins. A.D. and J.-Y.R. performed complementary studies in *D. melanogaster*. S.E. and K.Z. supervised the bioinformatics analyses. J.K. and P.B. conceived the project with K.Z. and supervised the experimental work. A.H., J.K., K.Z. and P.B. wrote the manuscript with help and comments from all co-authors.

## Supplemental Tables and Legends

**Supplemental Tables S1,3,4**

Provided as Excel files

**Supplementary Tables S2:**
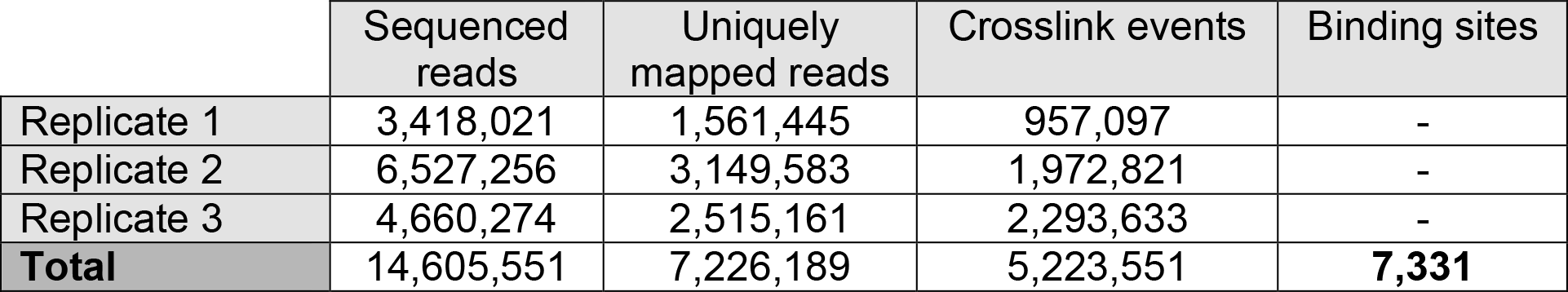
Summary of MKRN1 iCLIP experiments. iCLIP experiments with GFP-MKRN1 were performed in three independent replicates.

**Supplementary Table S5:**
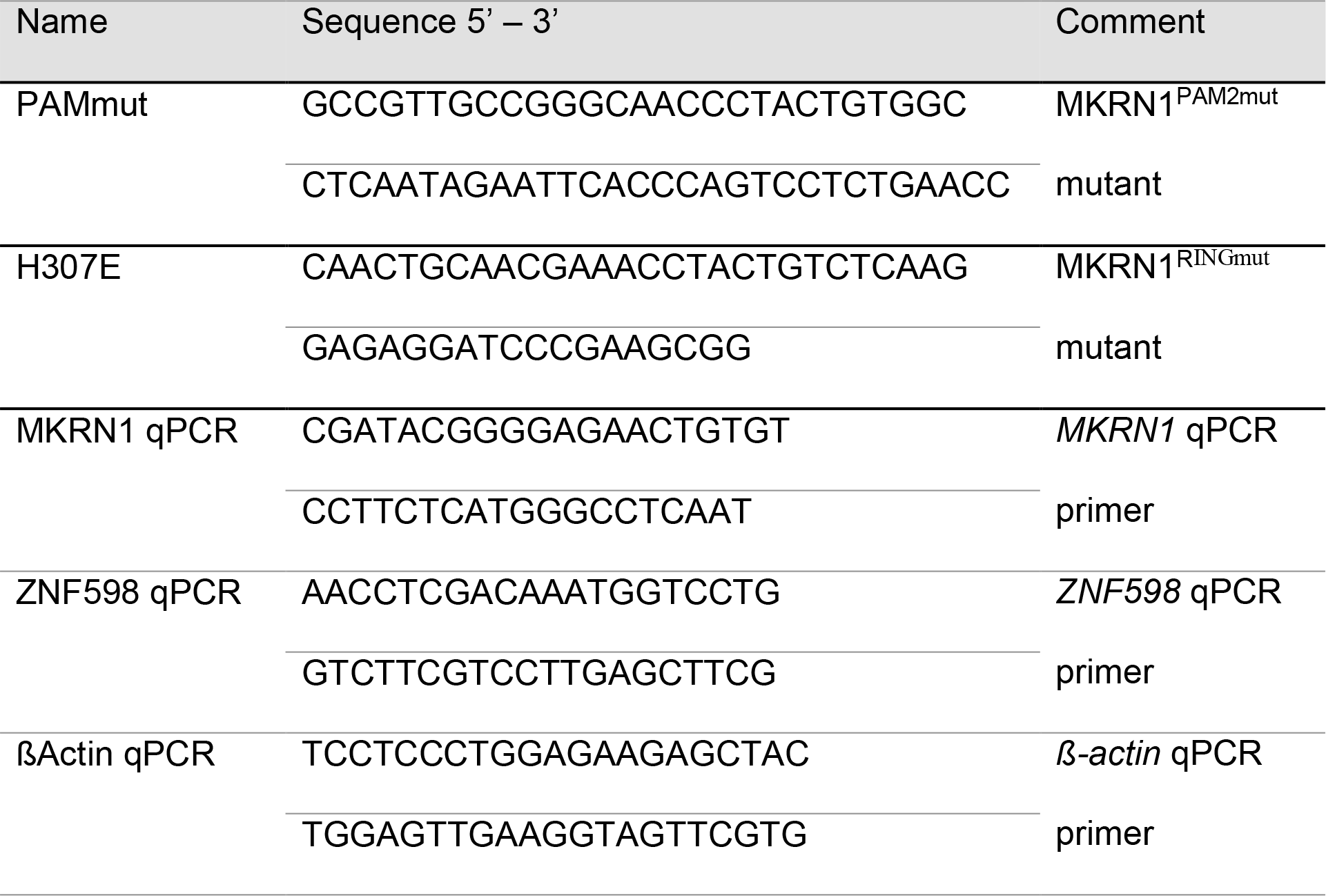
Primers used in this study

**Supplementary Table S6:**
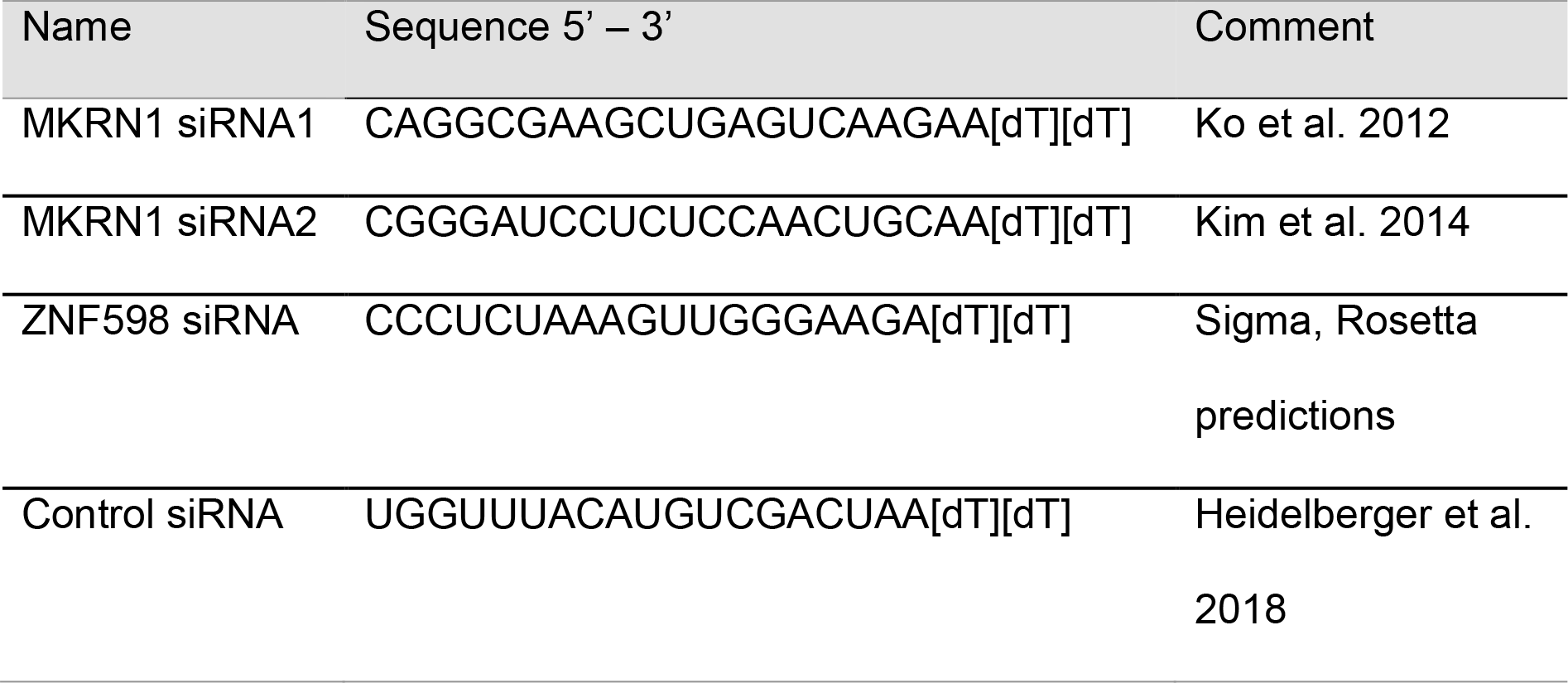
siRNAs used in this study

### Supplemental figure legends

**Supplementary Figure S1.**
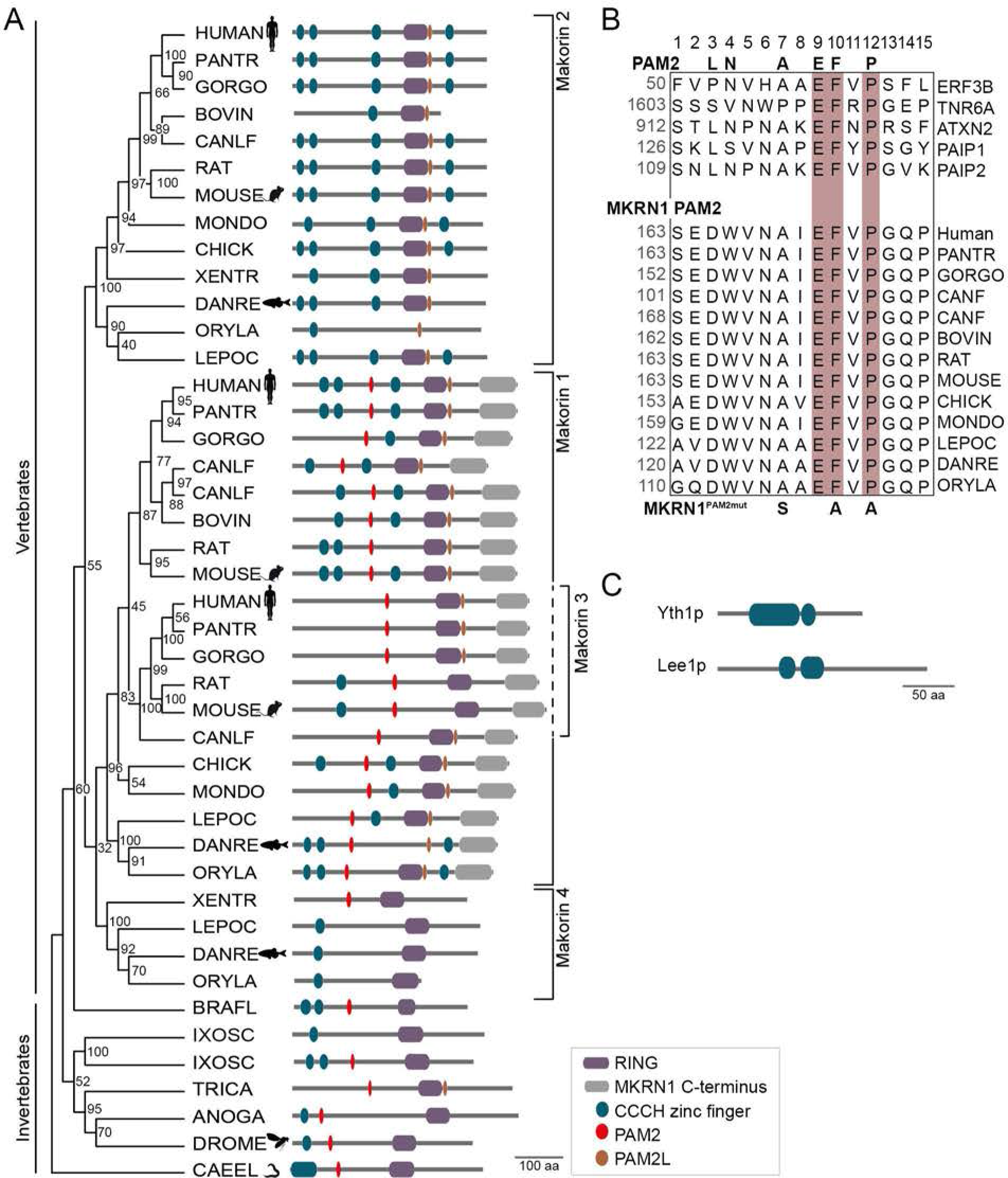
Maximum likelihood tree of Makorin orthologs with their protein domain architecture. (*A*) Maximum likelihood tree with 100 bootstrap replicates of selected vertebrate and invertebrate orthologs and *C. elegans* as an outgroup. Bootstrap values at each node indicate the number of replicates (out of 100) that support the local tree structure and thereby serve as confidence estimates. Protein schematics (drawn to scale) on the right depict protein domains corresponding to the following PFAM domains: RING-type zinc finger, PF13445; MKRN1 C-terminus, PF15815; CCCH zinc finger, PF15663, PF14608 and PF00642. PAM2 motifs, predicted to interact with the MLLE domain of PABP proteins (Kozlov et al. 2001) as well as the recently reported derivative PAM2L (Pohlmann et al. 2015), were added separately (see Material and methods). Abbreviated and full species names with corresponding UniProt identifiers in order of appearance: ANOGA, *Anopheles gambiae*, Q7QF83; BOVIN, *Bos taurus*, F1MF12, F6QQR5; BRAFL, *Branchiostoma floridae*, C3Y7M0; CAEEL, *Caenorhabditis elegans*, Q9N373; CANLF, *Canis lupus*, J9P921, E2RRA5, E2REH2, J9P9K3; DANRE, *Danio rerio*, Q4VBT5, Q9DFG8, A9C4A6; DROME, *Drosophila melanogaster*, Q9VP20; CHICKEN, *Gallus gallus*, Q9PTI4, F1NI93; GORGO, *Gorilla gorilla*, G3S6Y3, G3QDU4, G3RZ99; HUMAN, *Homo sapiens*, Q9UHC7, Q9H000, Q13064; IXOSC, *Ixodes scapularis*, B7QIJ9, B7Q4B2; LEPOC, *Lepisosteus oculatus*, W5NGW8, W5N9B2, W5LWJ1; MONDO, *Monodelphis domestica*, F6QPR3, F7F0I3; MOUSE, *Mus musculus*, Q9QXP6, Q9ERV1, Q60764; ORYLA, *Oryzias latipes*, H2MBR3, H2M1P4, H2LQG1; PANTR, *Pan troglodytes*, H2QVH8, H2QM29, H2Q915; RAT, *Rattus norvegicus*, A0A0G2QC40, Q5XI23, D3ZY41; XENTR, *Xenopus tropicalis*, Q6GLD9, B4F720. (*B*) The PAM2 motif in Makorin proteins from vertebrates (bottom, species abbreviations as in (*A*)) shows similarities to PAM2 in known PABP-interacting proteins from human (top, protein names given; first amino acid position for all PAM2 motifs indicated on the left in grey). The PAM2 consensus (Kozlov et al. 2001) is given above. Positions 9, 10 and 12 within the aligned regions that are highly consistent between all aligned proteins and important for PAM2 function (Kozlov et al. 2004) are highlighted in brown. Mutations that were introduced to abrogate the function of the PAM2 motif in human MKRN1 (MKRN1^PAM2mut^) are shown below. The corresponding UniProt identifiers are Q8IYD1, Q8NDV7, Q99700, Q9H074, Q9BPZ3 (known PABP-interacting proteins from human), Q9UHC7, H2QVH8, G3S6Y3, J9P921, E2RRA5, F1MF12, Q5XI23, Q9QXP6, Q9PTI4, F6QPR3, W5NGW8, Q4VBT5, H2MBR3 (Makorin orthologs from vertebrates). (*C*) The closest Makorin orthologs in *Saccharomyces cerevisiae* lack RING domain and PAM2 motif. Domain architecture of Yth1p and Lee1p, which were detected as closest orthologs by HaMStR-OneSeq (Ebersberger et al. 2014), but were not considered as orthologs due to low FAS scores (0,59 and 0,60, respectively). The annotated PFAM domains are CCCH zinc finger, PF15663, PF00642, PF16131.

**Supplementary Figure S2.**
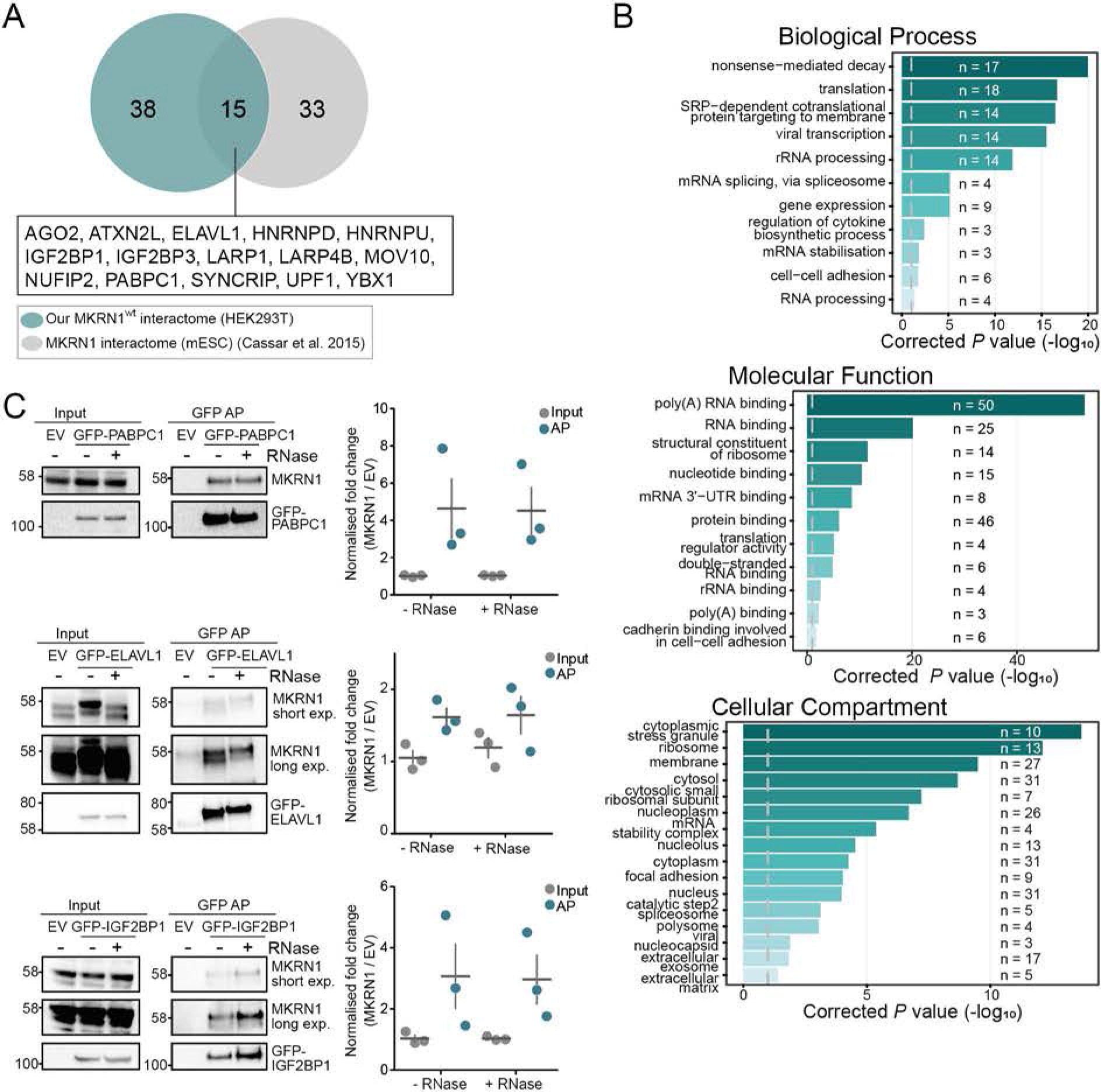
MKRN1 interacts with translational regulators and other RBPs. (*A*) Overlap of the 53 significant interaction partners of GFP-MKRN1^wt^ in human HEK293T cells with previously published interactors of MKRN1 in mouse embryonic stem cells (mESC) (Cassar et al. 2015). (*B*) GO terms enriched for the 53 MKRN1 interactors. *P* values (modified Fisher exact test, Benjamini-Hochberg correction) are depicted for all significant GO terms (corrected *P* value < 0.05) for Biological Process, Molecular Function and Cellular Compartment, together with the number of interactors associated with the respective term. (*C*) Reciprocal APs show that MKRN1 interacts with PABPC1, ELAVL1 and IGF2BP1 independently of RNA. AP with GFP-PABPC1, GFP-ELAVL1 and GFP-IGF2BP1 as baits were performed from HEK293T cells in the presence or absence of RNase A and T1. Bait proteins and endogenous MKRN1 were detected by Western blots (replicate 1). Different exposure times (exp.) for MKRN1 are shown for GFP-ELAVL1 and GFP-IGF2BP1 APs. Quantifications (fold changes of the MKRN1 signal over empty vector (EV)) of three replicates are shown on the right. Replicates 2 and 3, and uncropped gel images are shown in **Supplemental Fig. S9D-F**.

**Supplementary Figure S3.**
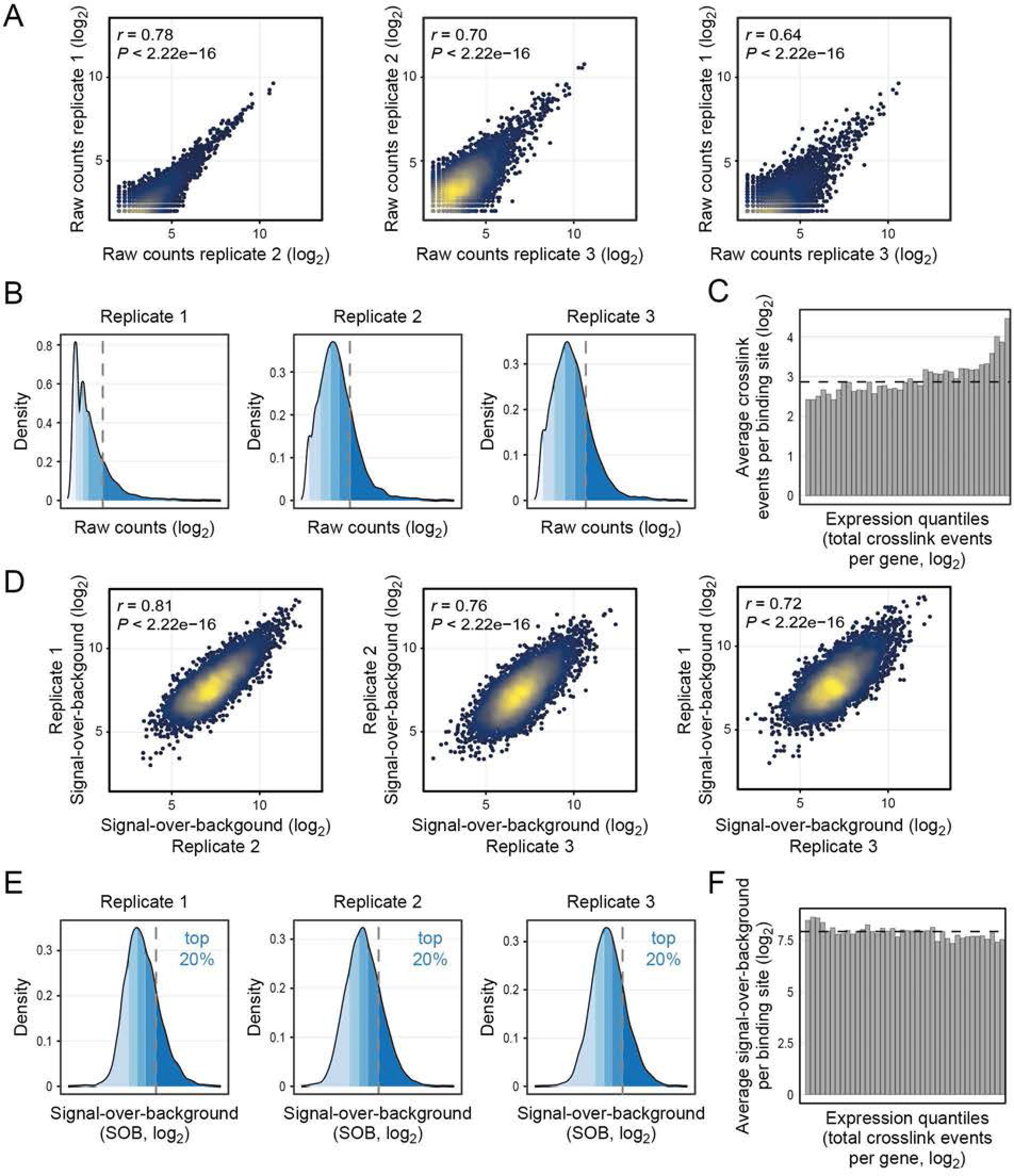
Signal-over-background transformation allows to estimate MKRN1 binding strength. (*A-C*) Raw iCLIP signal before signal-over-background transformation. (A) Scatter plots show pairwise comparisons of crosslink events per binding site in three replicate MKRN1 iCLIP experiments. Pearson correlation coefficients (*r*) and associated *P* values are given. (*B*) Density plots depict the distribution of crosslink events per binding site in the three replicate experiments. Shades of blue indicate 20% quantiles; top 20% of binding sites with highest counts are denoted by a dashed line. (*C*) Raw iCLIP counts are strongly influenced by the expression level of the underlying gene. MKRN1-bound genes were stratified into 50 bins with increasing expression (using the total number of MKRN1 crosslink events within the 3’ UTR as a proxy of a gene’s expression level). Shown is the average number of crosslink events per binding site for all binding sites in each bin. Dashed line denotes median across all bins. (*D-F*) Signal-over-background (SOB) values allow to correct for expression-level differences. (*D*) Pairwise comparison of SOB values for the three MKRN1 iCLIP replicate experiments. Scatter plots as in (*A*). (*E*) Distribution of SOB values in the three replicates. Density plots as in (*B*). Shades of blue indicate 20% quantiles. Dashed lines denote the top 20% MKRN1 binding sites with strongest binding that were used for the analyses in **Fig. 2B** and **Supplemental Fig. S4A**. (*F*) SOB values are independent of the expression level of the underlying gene. Average SOB values for all binding sites in each expression bin are shown as in (*C*).

**Supplementary Figure S4.**
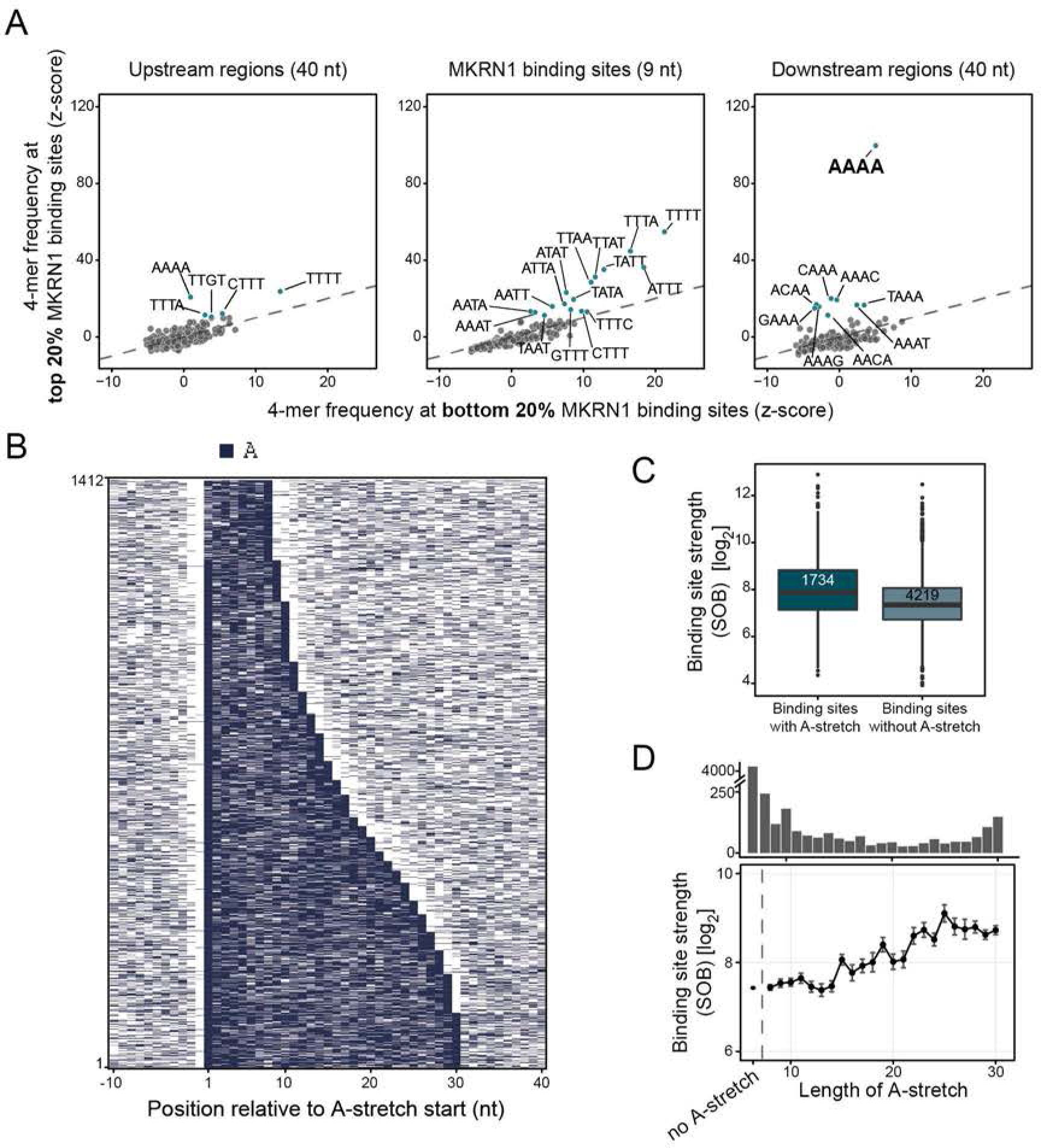
MKRN1 binds upstream of long A-stretches. (*A*) Binding sites with associated A-stretches show stronger MKRN1 binding. Boxplot compares the SOB values of MKRN1 binding sites in 3’ UTRs with and without associated A-stretches. Number of binding sites indicated inside box. (*B*) Heatmap representation of 1,412 non-overlapping A-stretches at MKRN1 binding sites, sorted by increasing length (8-30 nt). Only A’s are coloured. (*C*) MKRN1 binding site strength (signal-over-background, SOB) increases with length of associated A-stretch. Mean and standard deviation of MKRN1 binding sites associated with A-stretches of increasing length (x-axis). MKRN1 binding sites without associated A-stretches are shown for comparison on the left. Number of binding sites in each category indicated as barchart above. (*D*) The top 20% MKRN1 binding sites show a strong RNA binding preference for AAAA. Scatter plot compares the frequency of 4-mers within the 9-nt MKRN1 binding sites and flanking 40-nt windows for the top 20% and bottom 20% MKRN1 binding sites (according to SOB). 4-mer frequencies are displayed as z-scores based on background distribution from binding site permutations.

**Supplementary Figure S5.**
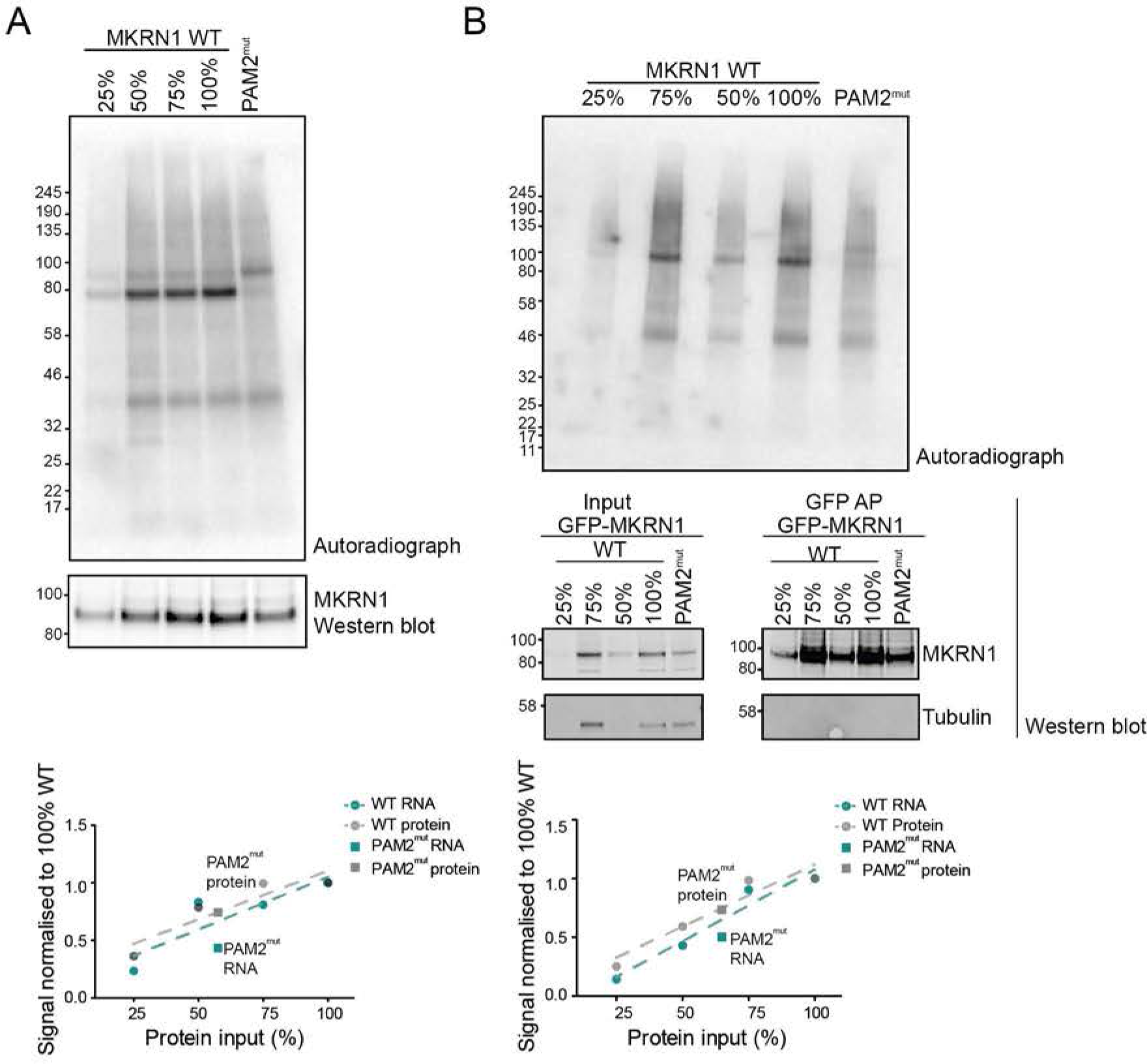
Interaction with PABP is required for MKRN1 RNA binding. (*A,B*) UV crosslinking experiments to measure the RNA binding capacity of GFP-MKRN1^wt^ and GFP-MKRN1^PAM2mut^. Autoradiographs (top) and Western blots (bottom) show GFP-MKRN1/RNA complexes and GFP-MKRN1 protein, respectively, in the eluates from replicates 2 (with 4SU and UV crosslinking at 365 nm) (*A*) and 3 (with conventional UV crosslinking at 254 nm) (*B*). For calibration, input samples for GFP-MKRN1^wt^ were diluted to 75%, 50% and 25% prior to GFP AP. Note that samples were loaded in different order in (*B*). Quantifications are given below. Uncropped gel images are shown in **Supplemental Fig. S10**.

**Supplementary Figure S6.**
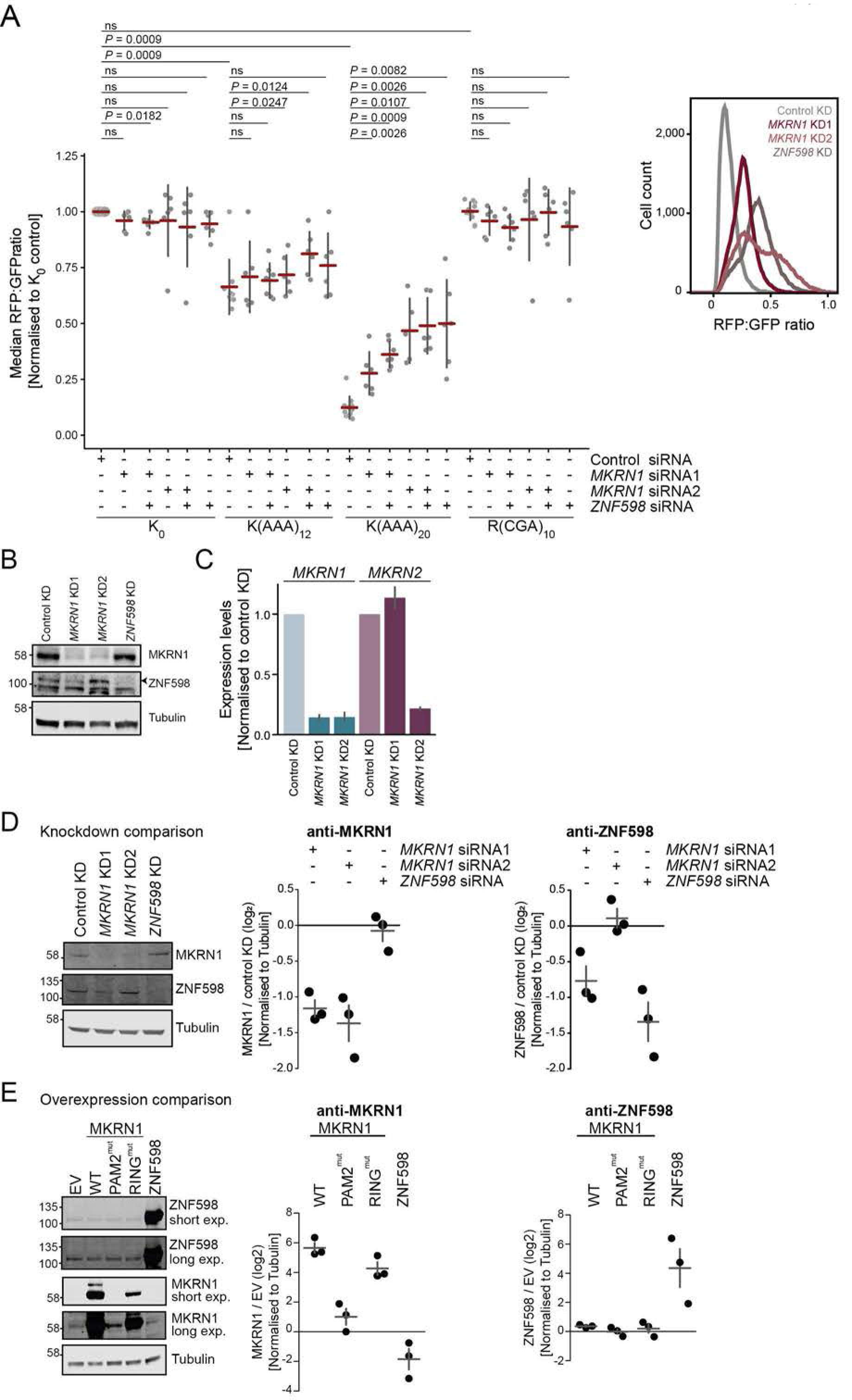
MKRN1 is required to stall ribosomes at K(AAA)_20_ in reporter assays. (*A*) Translation of dual fluorescence reporter plasmids was assessed by flow cytometry upon *MKRN1* and/or *ZNF598* KD. Median RFP:GFP ratios (normalised to K_0_ in control KD) are shown for the reporter plasmids K_0_, K(AAA)_12_, K(AAA)_20_, and R(CGA)_10_. Error bars represent standard deviation of the mean (s.d.m., n ≥ 6 replicates; paired two-tailed t-test, Benjamini-Hochberg correction). Density plot of median RFP:GFP ratios of one replicate experiment with K(AAA)_20_ with control or *MKRN1* KD (two independent siRNAs, KD1 and KD2) or *ZNF598* is shown on the right. KDs of *MKRN1* and *ZNF598* were assessed by Western blot (n = 3 replicates). Black arrowhead indicates ZNF598. Replicates 2 and 3, and uncropped gel images are shown in **Supplemental Fig. S11A,B**. (*C*) *MKRN1* KD2 also reduces *MKRN2* levels. *MKRN1* KD1 and KD2 were performed for 72 h. Expression levels of *MKRN1* and *MKRN2* were assessed in relation to *ß-actin* levels by qPCR in *MKRN1* KD (siRNA 1 and 2) and control KD. Error bars indicate s.d.m. (n = 2 replicates). (*D,E*) Cross-regulation of MKRN1 and ZNF598. (*D*) *MKRN1* KD1 reduces endogenous ZNF598 protein levels. Effect of *MKRN1* KD (KD1, siRNA 1 and KD2, siRNA 2) and *ZNF598* KD for 72 h was assessed by Western blot for endogenous MKRN1 and ZNF598. Quantifications depict MKRN1 or ZNF598 expression levels in *MKRN1* or *ZNF598* KD over control KD condition, normalised to tubulin levels (n = 3 replicates). Replicates 2 and 3, and uncropped gel images are shown in **Supplemental Fig. S11C,D.**(*E*) *ZNF598* overexpression reduces MKRN1 protein levels. Effect of *ZNF598* and *MKRN1* (wt and mutants) overexpression was tested after 48 h. Quantification as in (*D*). Uncropped gel images for all replicates are shown in **Supplemental Fig. S11E,F**.

**Supplementary Figure S7.**
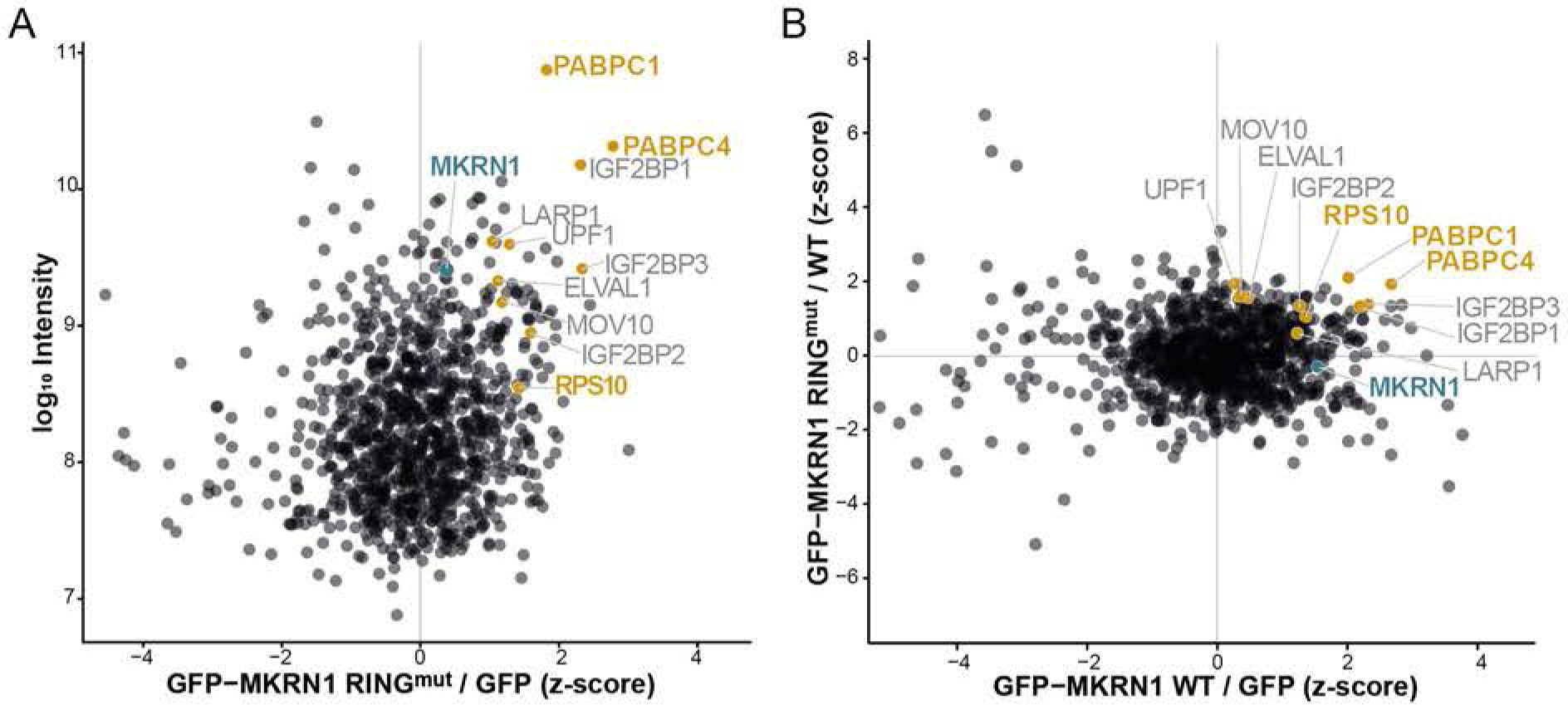
Interactome of GFP-MKRN1^RINGmut^ reveals putative ubiquitylation substrates. Experiments were performed using SILAC-based MS. Asymmetrical z-scores of combined SILAC ratios (n = 3 replicates) are shown. Proteins are detected in at least two out of three replicates. (*A*) Protein interactome of GFP-MKRN1^RINGmut^ in HEK293T cells analysed by quantitative mass spectrometry. Combined SILAC ratios (n = 3 replicates) after z-score normalisation are plotted against log_10_-transformed intensities. 1,097 protein groups were quantified in at least two out of three replicates (**Supplemental Table S1**). MKRN1 and interesting candidate ubiquitylation targets are highlighted. (*B*) Quantitative comparison of the interactome of GFP-MKRN1^wt^ and GFP-MKRN1^RINGmut^ shows that potential ubiquitylation candidates identified in (*A*) are enriched in GFP-MKRN1^RINGmut^ over GFP-MKRN1^wt^. Comparison reveals 137 proteins to be significantly enriched (MKRN1^RINGmut^ over MKRN1^wt^ with FDR < 5% and MKRN1^wt^/GFP z-score > 1). Combined ratios of three replicates are shown in a scatter plot. Only proteins detected in at least two out of three replicates are shown. Highlighting as in (*A*).

**Supplementary Figure S8.**
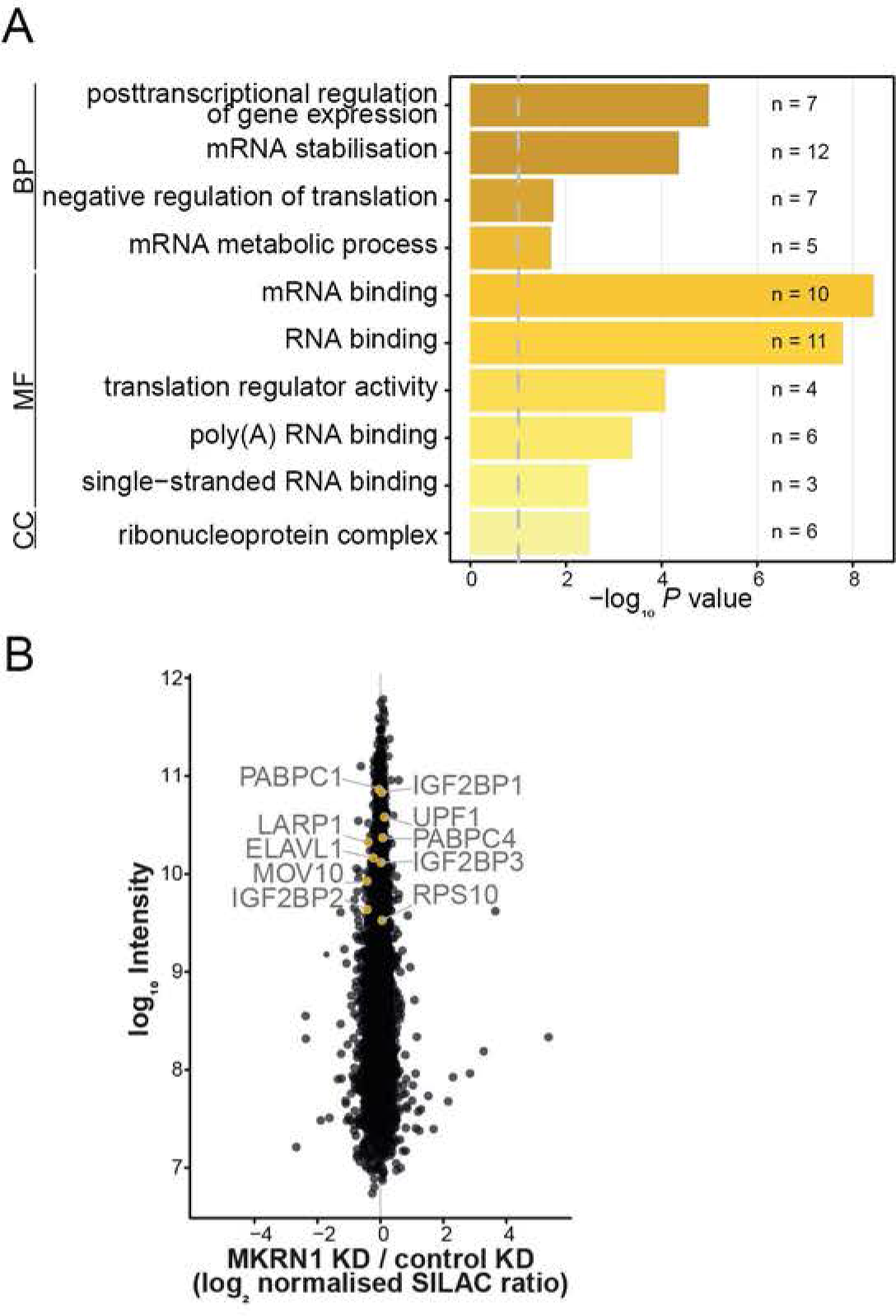
GO term analysis of MKRN1 ubiquitylation targets and proteome analysis upon *MKRN1* KD. (*A*) GO terms enriched for the 21 MKRN1 ubiquitylation targets. Corrected *P* values (modified Fisher exact test, Benjamini-Hochberg correction) are depicted for all significant GO terms (corrected *P* value < 0.05) for Biological Process (BP), Molecular Function (MF) and Cellular Compartment (CC), together with the number of ubiquitylation targets associated with the respective term. (*B*) Proteome analysis of *MKRN1* KD in HEK293T cells analysed by quantitative mass spectrometry. Log_2_-transformed, combined normalised SILAC ratios (n = 3 replicates) are plotted against log_10_-transformed intensities. 6,425 protein groups were quantified in at least one out of three replicate experiments (**Supplemental Table S4**). Selected ubiquitylation targets of MKRN1 are highlighted.

**Supplementary Figure S9.**
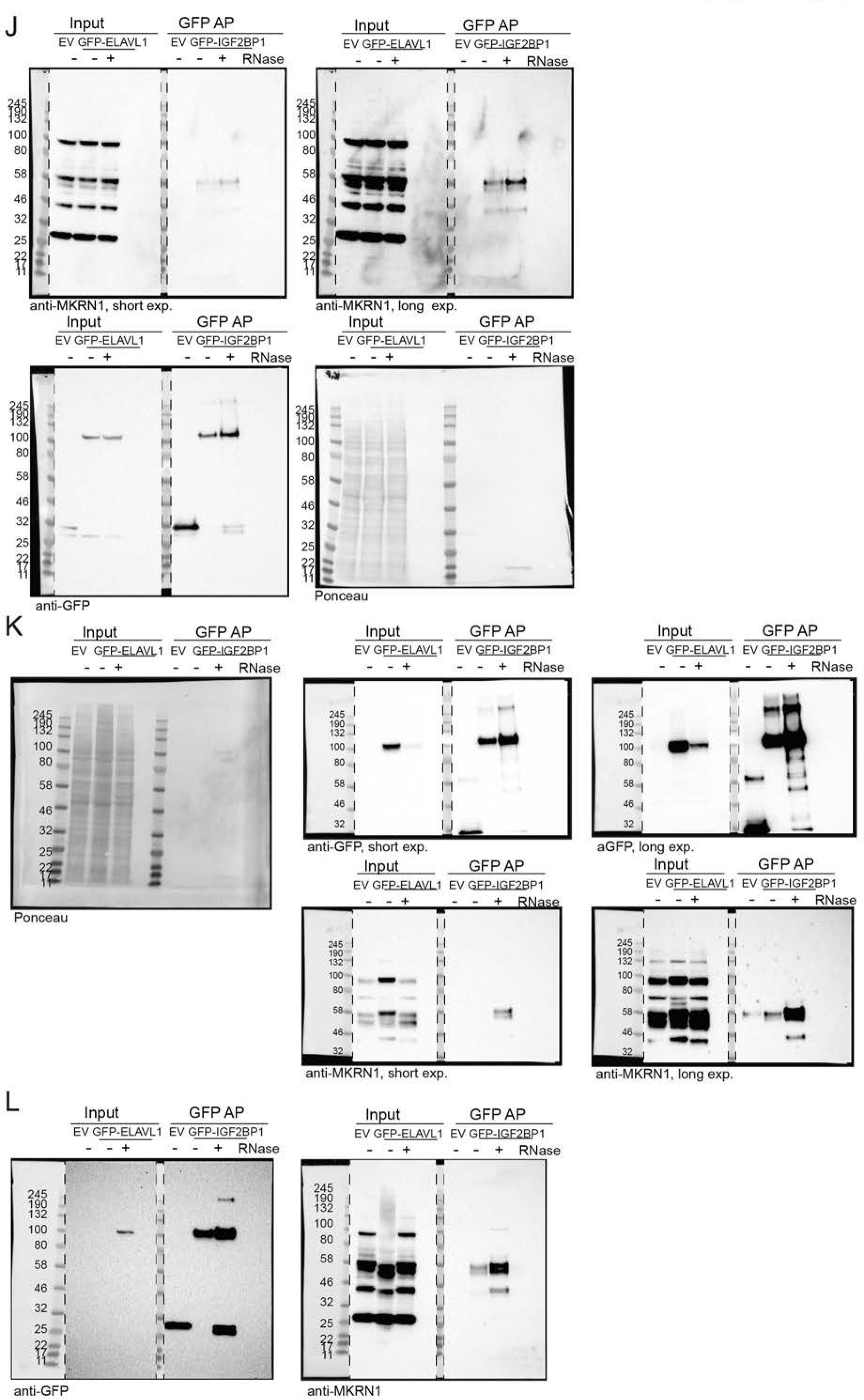
Images of full membranes and different exposure times (exp.) for Western blot analyses in **Fig. 1C** and **Supplemental Fig. S2C** in the presence or absence of RNase A and T1. (*A-C*) PABP interacts with MKRN1^wt^ and MKRN1^RINGmut^ but not MKRN1^PAM2mut^. Western blot analysis was performed with antibodies against PABPC1/3 and GFP. Images of full membranes and different exposure (exp.) times for both antibodies are shown for replicate 1 (*A*) which is presented in **Fig. 1C**, as well as replicates 2 (*B*) and 3 (*C*). Black and blue arrowheads indicate GFP-MKRN1 and PABPC1/3, respectively. (*D-F*) Endogenous MKRN1 interacts with GFP-PABPC1 independent of RNA. Western blot analysis was performed with antibodies against MKRN1 and GFP. Images of full membranes and different exposure times for both antibodies are shown for replicate 1 (*D*) which is presented in **Supplemental Fig. S2C**, as well as replicates 2 (*E*) and 3 (*F*). Black and blue arrowheads indicate MKRN1 and GFP-PABPC1, replicates. (*G-I*) Endogenous MKRN1 interacts with GFP-ELAVL1 independent of RNA. Western blot analysis was performed with antibodies against MKRN1 and GFP. Images of full membranes and different exposure times for both antibodies are shown for replicate 1 (*G*) which is presented in **Supplemental Fig. S2C**, as well as replicates 2 (*H*) and 3 (*I*). (*J-L*) Endogenous MKRN1 interacts with GFP-IGF2BP1 independent of RNA. Western blot analysis was performed with antibodies against MKRN1 and GFP. Images of full membranes and different exposure times for both antibodies are shown for replicate 1(*J*) which is presented in **Supplemental Fig. S2C**, as well as replicates 2 (*K*) and 3 (*L*).

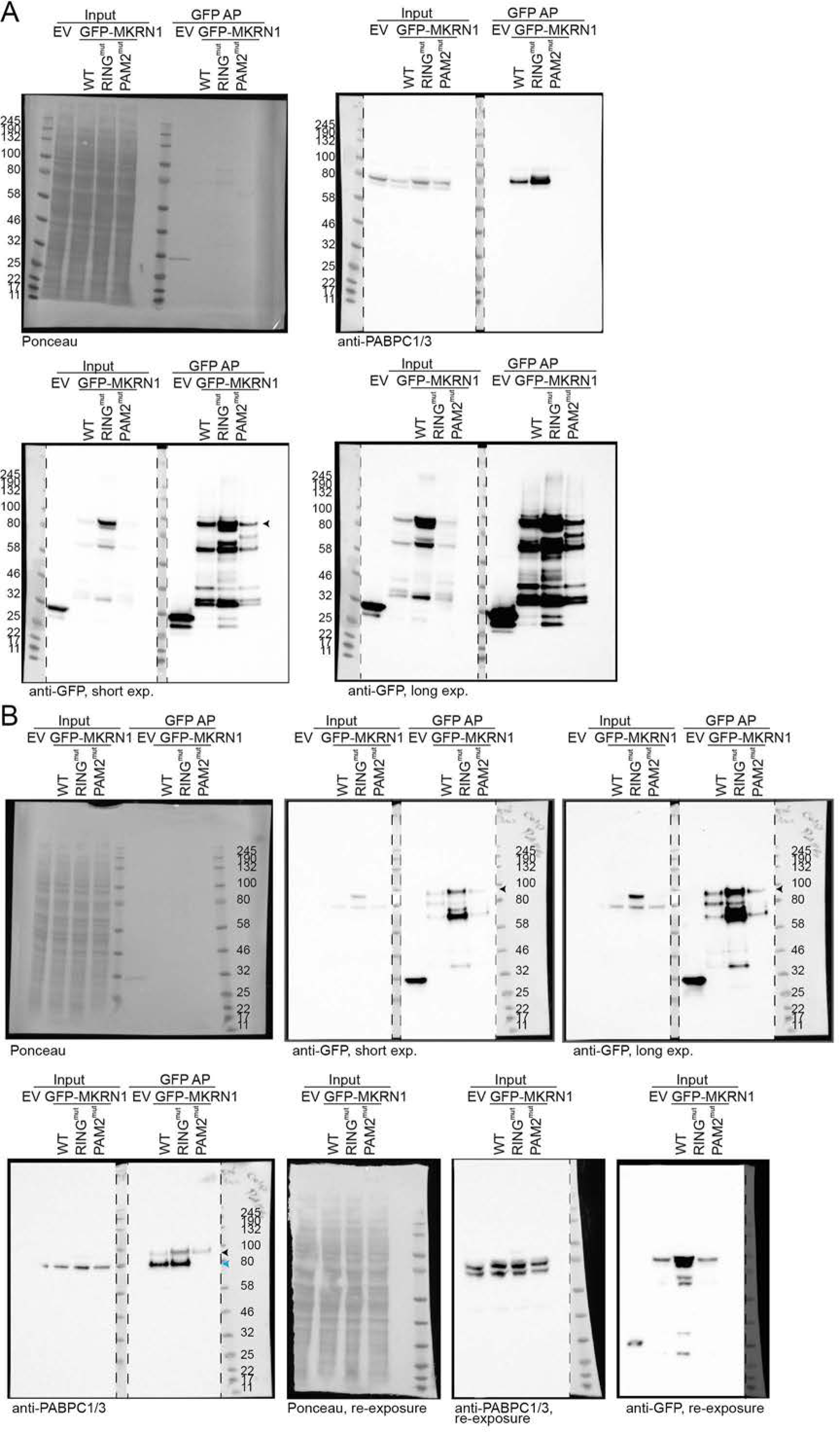

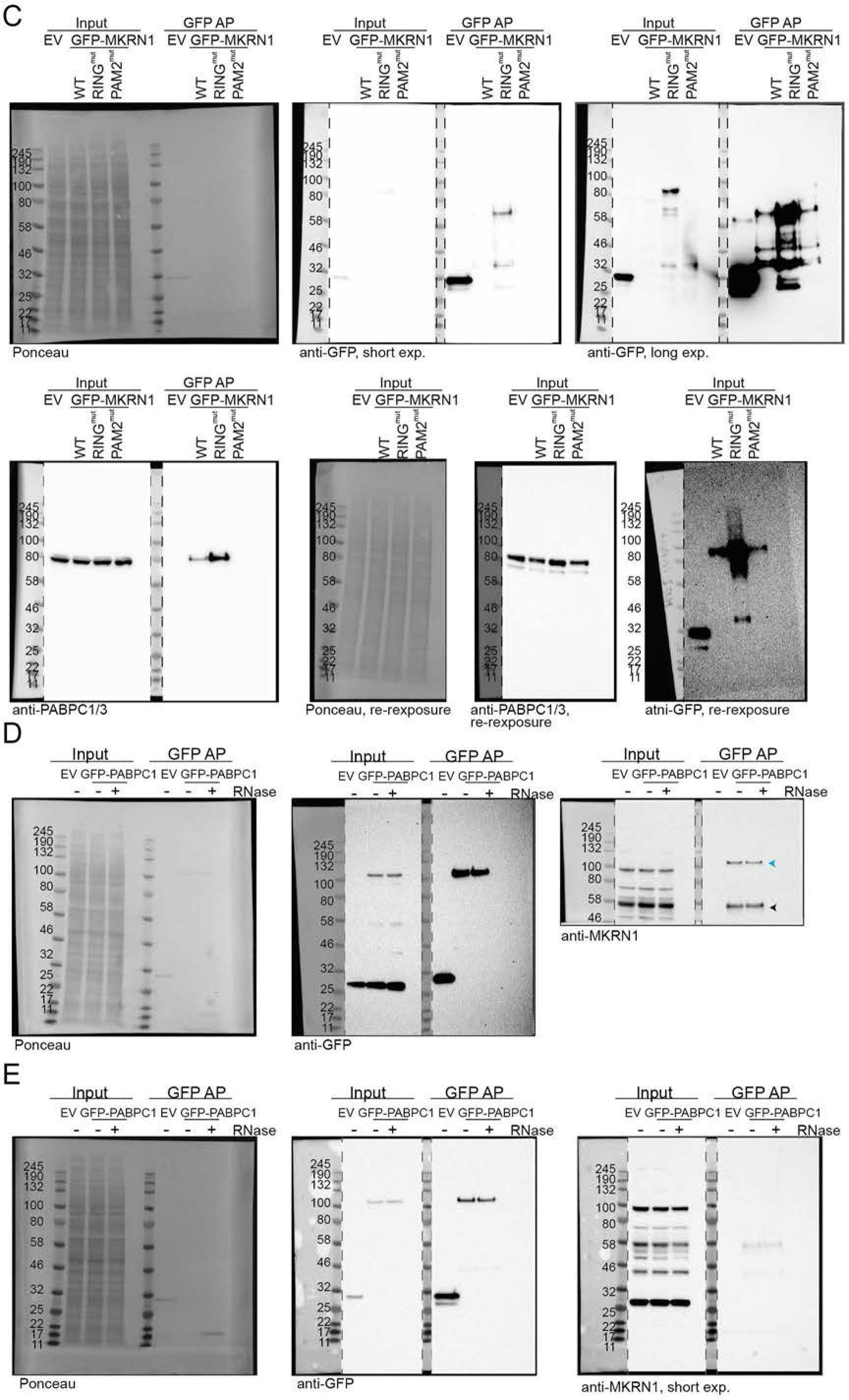

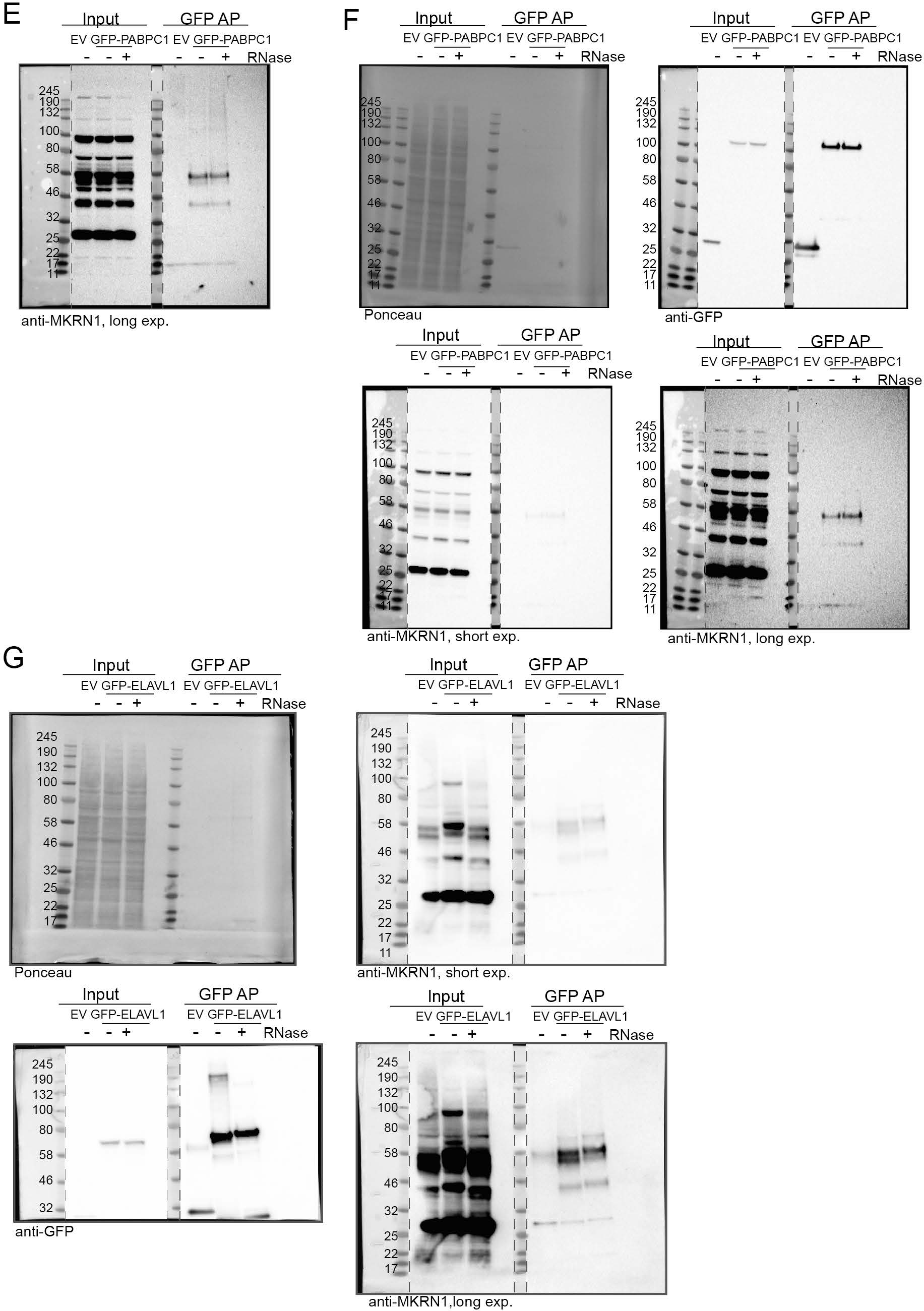

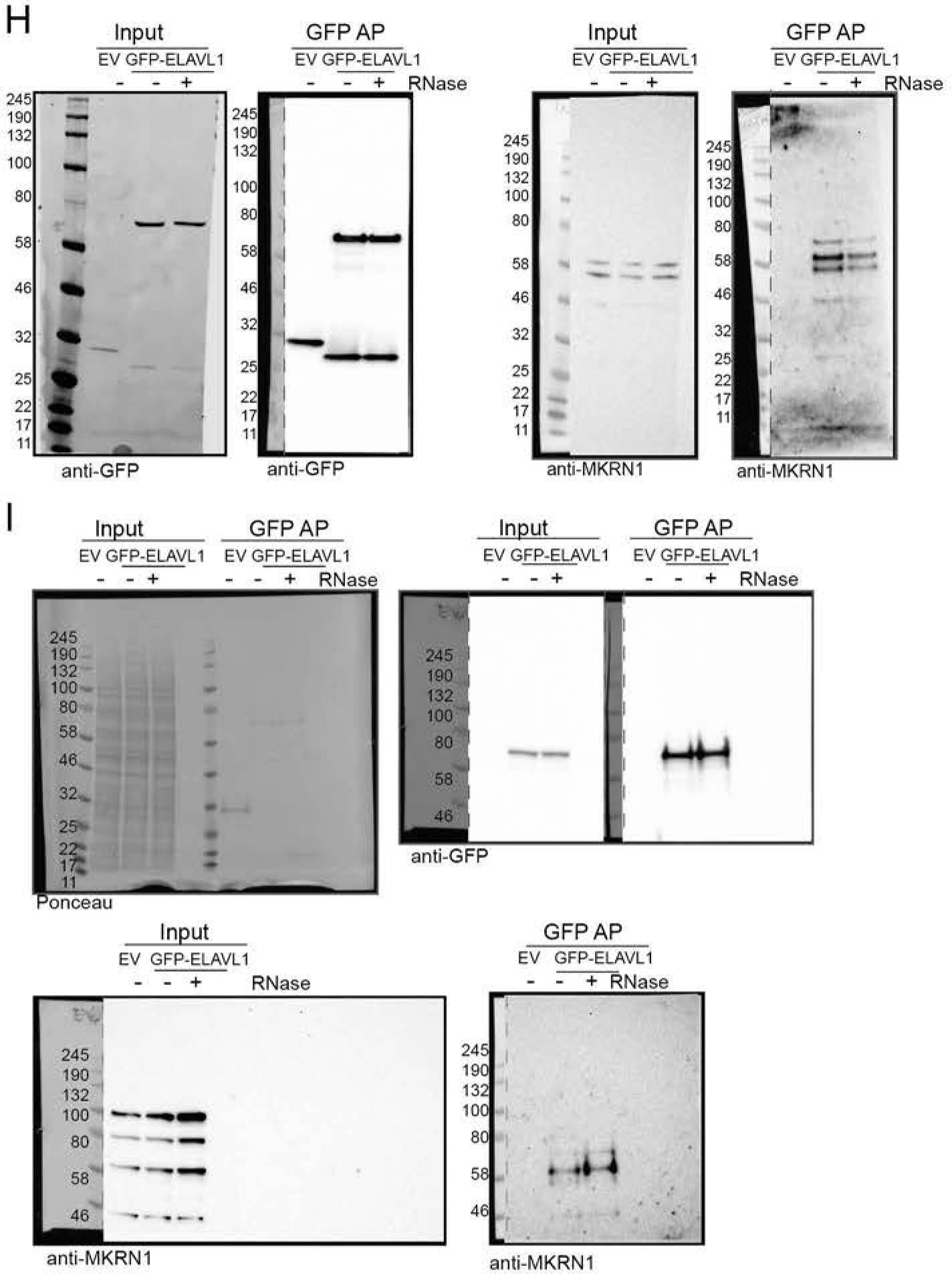

**Supplementary Figure S10.**
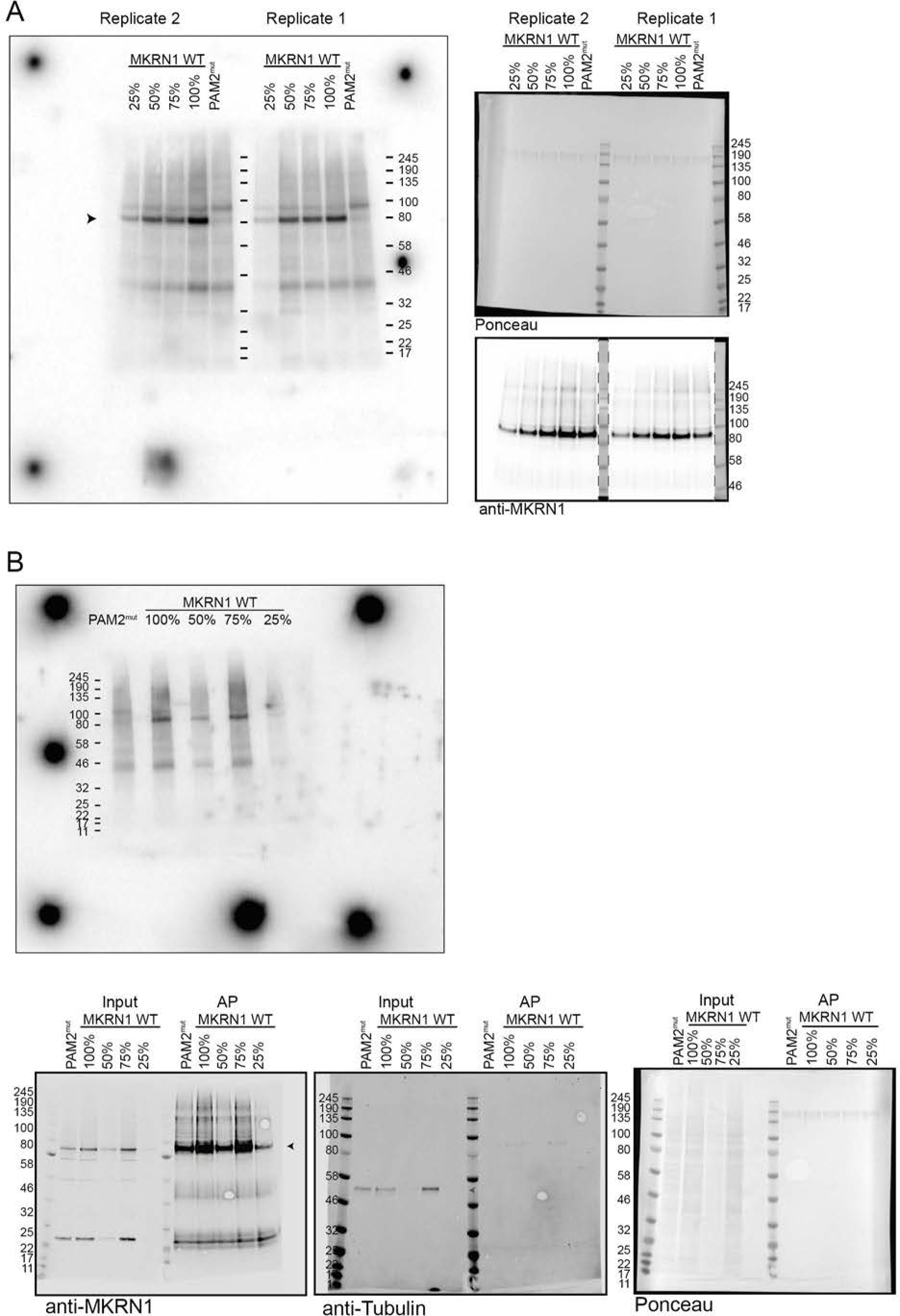
Images of full membranes of autoradiographs and Western blot analyses in **Fig. 3D**(replicate 1) and **Supplemental Fig. S5**(replicates 2 and 3). UV crosslinking experiments to measure the RNA binding capacity of GFP-MKRN1^wt^ and GFP-MKRN1^PAM2mut^. Autoradiographs (*A*, left; *B*, top) and Western blots (*A*, right; *B*, bottom) show GFP-MKRN1/RNA complexes and GFP-MKRN1 protein, respectively, in the eluates from replicates 1 and 2 (with 4SU and UV crosslinking at 365 nm) (*A*) and 3 (with conventional UV crosslinking at 254 nm) (*B*). (*B*) Images of full membranes of Western blot analyses with both antibodies are shown for replicate 3 (*B*).

**Supplementary Figure S11.**
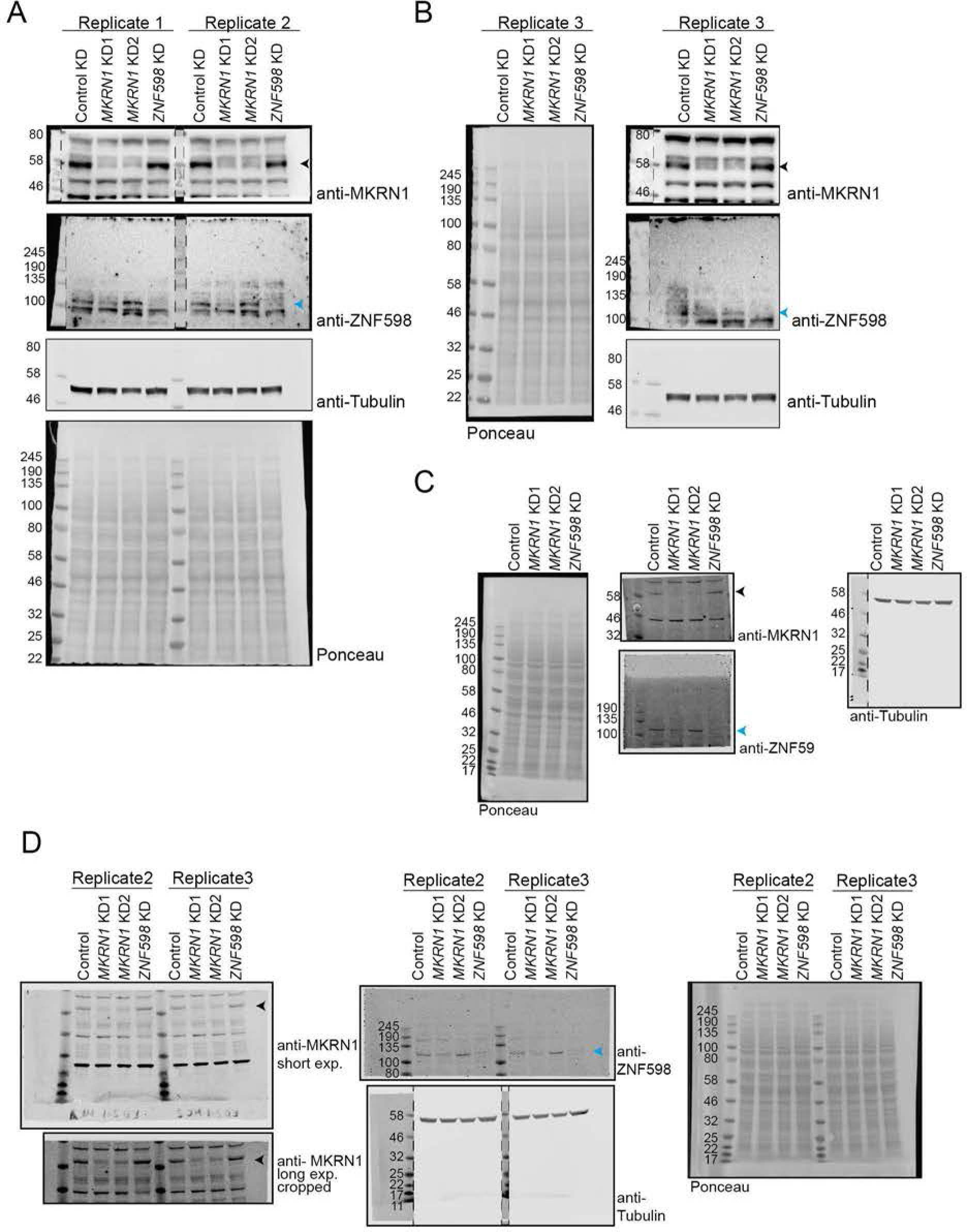
Images of full membranes and different exposure (exp.) times for Western blot analyses in **Supplemental Fig. S6B,D,E.**(*A,B*) KDs of *MKRN1* and *ZNF598* assessed by Western blot (n = 3 replicates) from **Supplemental Fig. S6B**. Western blot analysis was performed with antibodies against MKRN1, ZNF598, and tubulin. Black and blue arrowheads indicate MKRN1 (53 kDa) and ZNF598 (99 kDa), respectively. Uncropped gel images of replicates 1 and 2 (*A*) and 3 (*B*). (*C,D*) Images of full membranes are shown for cross-regulation between *MKRN1* and *ZNF598* KD from **Supplemental Fig. S6D**. *MKRN1* KD1 reduces endogenous ZNF598 protein levels. Western blot analysis was performed with antibodies against MKRN1, ZNF598, and tubulin. Coloured arrowheads as in (*A*). Uncropped gel images of replicate 1 (*C*) and replicates 2 and 3 (*D*). (*E,F*) Images of full membranes are shown for cross-regulation of *MKRN1* and *ZNF598* overexpression (OE) from **Supplemental Fig. S6E**. *ZNF598* OE reduces MKRN1 protein levels. Western blot analysis was performed with antibodies against MKRN1, ZNF598, and tubulin. Black arrowheads indicate MKRN1. Images of full membranes and different exposure times (exp.) for both antibodies are shown for replicates 1, 2 (*E*), and 3 (*F*). Note the opposite order of replicates 1 and 2 (2 left, 1 right) in (*E*).

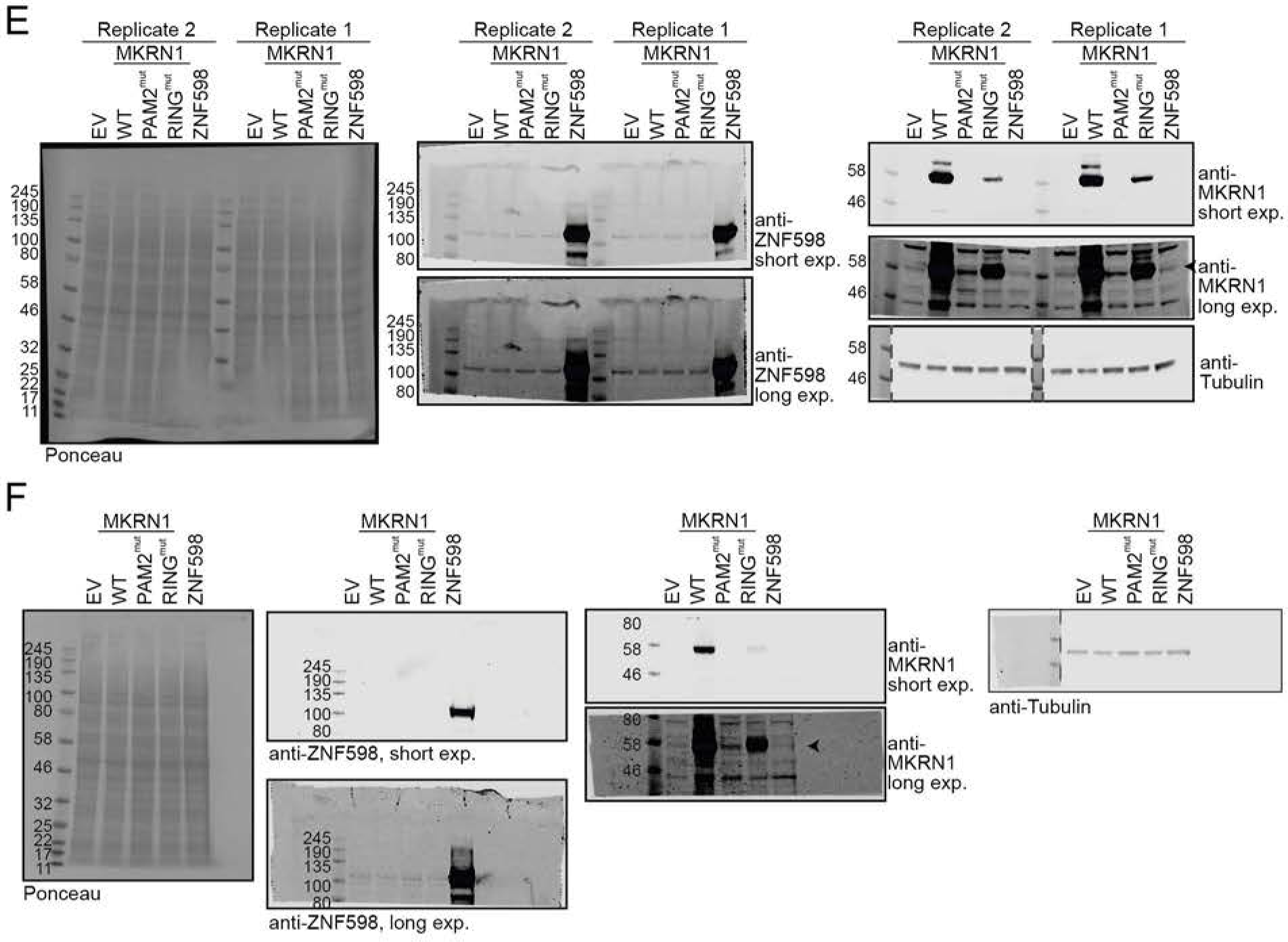

## Abbreviations

Adenosine: (A)
Makorin Ring Finger Protein 1: (MKRN1)
ribosome-associated quality control: (RQC)
poly(A)-binding protein: (PABP)
mass spectrometry: (MS)
affinity purification: (AP)
stable isotope labelling with amino acids in cell culture: (SIALC)
false discovery rate: (FDR)
Gene Ontology: (GO)
Biological Process: (BP)
Molecular Function: (MF)
mRNA ribonucleoprotein particle: (mRNP)
PABP-interacting motif: (PAM2)
MKRN1 variant with point mutations in the PAM2 motif: (GFP-MKRN1^PAM2mut^)
MKRN1 variant with point mutation in RING domain: (GFP-MKRN1^RINGmut^)
individual-nucleotide resolution UV crosslinking and immunoprecipitation: (iCLIP)
signal-over-background: (SOB)
nucleotides: (nt)
4-thiouridine: (4SU)
Pearson correlation coefficients: (r)
A-rich stretches: (A-stretches)
lysine: (K)
knock down: (KD)
RNA recognition motif: (RRM)
Polyethylenimine: (PEI)
modified RIPA: (mRIPA)
N-ethylmaleimide: (NEM)
dithiothreitol: (DTT)
chloroacetamide: (CAA)
higher-energy collisional dissociation: (HCD)
glycine-glycine: (GlyGly)
strong-cation exchange chromatography: (SCX)

